# Transcriptome profiling of the ventral pallidum reveals a role for pallido-thalamic neurons in cocaine reward

**DOI:** 10.1101/2021.10.20.465105

**Authors:** Michel Engeln, Megan E. Fox, Ramesh Chandra, Eric Y. Choi, Hyungwoo Nam, Houman Qadir, Shavin S. Thomas, Victoria M. Rhodes, Makeda D. Turner, Rae J. Herman, Cali A. Calarco, Mary Kay Lobo

**Affiliations:** Department of Anatomy and Neurobiology, University of Maryland School of Medicine, Baltimore, MD, USA; University of Bordeaux, CNRS, INCIA, UMR 5287, F-33000 Bordeaux, France; Department of Anesthesiology & Perioperative Medicine, Penn State College of Medicine, Hershey, PA, USA

**Author notes:** Address Correspondence to Mary Kay Lobo, 20 Penn St, HSFII Building, Rm 265, Baltimore, MD 21201, USA, Or Michel Engeln, Institut de Neuroscience Cognitives et Intégratives d’Aquitaine - UMR 5287, 146 rue Léo Saignat, Zone Nord - Bat 1A - 3e étage, 33076 Bordeaux cedex, France. Funding: This work was funded by NIH grants R01MH106500, R01DA038613, R01DA047843 and Israel-US Binational Science Foundation 201725 (to MKL), K99DA050575 (to MEF), F31DA052967 (to EC), T32DK098107 and F32DA052966 (to CAC).

## Abstract

Psychostimulant exposure alters the activity of ventral pallidum (VP) projection-neurons. However, the molecular underpinnings of these circuit dysfunctions are unclear. We used RNA-sequencing to reveal alterations in the transcriptional landscape of the VP that are induced by cocaine self-administration in mice. We then probed gene expression in select VP neuronal subpopulations to isolate a circuit associated with cocaine intake. Finally, we used both overexpression and CRISPR-mediated knockdown to test the role of a gene target on cocaine-mediated behaviors as well as dendritic spine density. Our results showed that a large proportion (55%) of genes associated with structural plasticity were changed 24 hours following cocaine intake. Among them, the transcription factor Nr4a1 (Nuclear receptor subfamily 4, group A, member 1, or Nur77) showed high expression levels. We found that the VP to mediodorsal thalamus (VP→MDT) projection neurons specifically were recapitulating this increase in Nr4a1 expression. Overexpressing Nr4a1 in VP→MDT neurons enhanced drug-seeking and drug-induced reinstatement, while Nr4a1 knock down prevented self-administration acquisition and subsequent cocaine-mediated behaviors. Moreover, we showed that Nr4a1 negatively regulated spine dynamics in this specific cell subpopulation. Together, our study identifies for the first time the transcriptional mechanisms occurring in VP in drug exposure. Our study provides further understanding on the role of Nr4a1 in cocaine-related behaviors and identifies the crucial role of the VP→MDT circuit in drug intake and relapse-like behaviors.

## Introduction

Originally associated with motor functions, the ventral pallidum (VP) has gained attention for its role in motivation and reward seeking behaviors ^1-5^. Receiving inputs from both nucleus accumbens (NAc) projection neuron subtypes ^6, 7^ as well as direct mesolimbic dopamine projections ^8^, the VP holds a central place in the reward circuitry. Early work investigating its potential involvement in addiction quickly highlighted its essential role in cocaine-related behaviors ^8-10^. With the development of new tools, studies dissecting the precise role of the VP in reward revealed a heterogenous brain region composed of diverse cell subtypes and sub- circuits ^1, 11, 12^. Ventral pallidum neurons have different roles in reward processing based on their cellular identity (e.g., glutamateric vs. GABAergic, the expression of calbindin vs. parvalbumin, etc.) or output structure ^1, 3, 11, 13^, and undergo specific plastic changes following drug exposure ^2, 7, 12^. For instance, recent studies demonstrated opposing functions for VP projections to the ventral tegmental area (VTA) and the lateral habenula (LHb), either promoting or reducing reward-related behaviors respectively ^3, 11^. However, while a growing number of studies show changes in neuronal activity following drug exposure, data about molecular adaptations supporting these plastic changes in the VP is lacking. Investigating these transcriptomic adaptations could indeed reveal new molecular and neuronal pathways associated with drug intake.

In this study, we interrogated transcriptomic changes in the VP of mice following cocaine self-administration. Guided by bioinformatics analysis, we identified the transcription factor Nr4a1 (Nuclear receptor subfamily 4, group A, member 1, or *Nur77*) as a potential mediator of cocaine-induced structural plasticity. Subsequent circuit-specific gene expression evaluation revealed high Nr4a1 expression levels, as well as several of its target genes, specifically in VP neurons projecting to the MDT (VP→MDT) following cocaine intake. Manipulating Nr4a1 expression in this circuit bidirectionally altered cocaine-mediated behaviors as well as dendritic spine density, confirming both the role of Nr4a1 in drug-induced structural plasticity and the critical role of the VP→MDT circuit in cocaine reward.

## Material and Methods

### Animals

Studies were conducted in accordance with guidelines set up by the Institutional Animal Care and Use Committees at University of Maryland School of Medicine. All animals were given food and water ad libitum during the study with the exception of water training. Adult male and female wild-type C57BL/6 mice (University of Maryland Veterinary Resources) and male RiboTag (RT)^+/+^ mice (Rpl22^tm1.1Psam/J^) on a C57Bl/6 background ^14^ were used. To avoid any damage on catheters, during the entire behavioral study mice were pair-housed in cages with perforated plexiglass dividers to allow for sensory, but not physical, interaction ^15^. Mice were 7 to 8 weeks old at the beginning of the experiments.

### Intravenous Cocaine Self-administration

Self-administration experiments involving viral manipulation were conducted in two independent cohorts of animals, at least. Data were then pooled. Experimenter were not blind to the drug/virus conditions.

Intravenous cocaine self-administration experiments were conducted as described previously ^16, 17^. Briefly, in order to habituate mice to the nosepokes and facilitate operant responding to obtain a reward, water-deprived mice were trained to self-administer water (4 days, 2 sessions of 30 min/day) under an FR1 schedule ^16-19^. Active responses, under a fixed ratio 1 (FR1) schedule, elicited the delivery of a 10-μL water drop associated with a light-cue both delivered in the nosepoke hole. Reward delivery was followed by a 10-s timeout period and the house-light was turned off during this time. Following training, animals with stable operant responding were anesthetized with ketamine (100 mg/kg) and xylazine (16 mg/kg) and implanted with long-term indwelling jugular catheters (Plastics One). Mice were flushed daily with heparin 50 IU/mL (40%, in saline) and 2.27% Baytril (Bayer; 20%). After 5 days of recovery, mice underwent 10 days of saline or cocaine self-administration under an FR1 schedule (0.5 mg/kg/infusion in saline; 2 hours/day; randomly assigned to saline or cocaine group). Responses in the active nosepoke elicited a 10-μl cocaine (or saline) infusion and a 10-s illumination of the active nosepoke light. During this 10-s time out period, nosepokes in the active port were recorded but did not trigger additional infusions. Responses on the inactive nosepoke were also recorded but had no programmed consequences.

To avoid the effect of anesthetic on behavioral and molecular measurements during self-administration experiments, catheter patency was deduced from flawless daily heparin flushing and was inferred from daily increase in cocaine intake and from psychomotor activation at the end of each session in mice taking drug ^15^. Only mice showing these criteria were used in data analyses.

### Cocaine seeking, Extinction and Drug-induced reinstatement

Twenty-four hours after the last FR1 session, a 1-h seeking test was performed under extinction conditions (i.e., a response resulted in cue presentation but no drug delivery).

Following the seeking test, mice were subjected to additional extinction sessions consisting of 5 distinct 1-h sessions separated by 5-min intervals during which animals were placed back in their home cage. Similar to the seeking test, during extinction sessions each response on the active nosepoke resulted in cue presentation but no drug delivery. On the following day, the extinguished mice were injected with cocaine (7.5 mg/kg in saline, ip.) or saline 5 min before a cocaine induced reinstatement test during which each response resulted in cue presentation but no drug delivery ^15, 16, 20-22^.

### Stereotaxic surgery and viral vectors

Under isoflurane anesthesia, mice were bilaterally injected with viral vectors at the following coordinates (from bregma): ventral pallidum (+/-10° angle from dorsal skull surface, anterior/posterior: AP +0.9, medial/lateral: ML ±2.2, dorsal/ventral: DV -5.3), mediodorsal thalamus (10° angle, AP: -0.8, ML: ±1.2, DV: -3.7), ventral tegmental area (7° angle, AP: -3.2, ML: ±1, DV: -4.6), lateral habenula (10° angle, AP: -1.2, ML: ±0.7, DV: -3.1). Circuit-specific viral expression was achieved using the retrograde properties of adeno-associated virus (AAV) serotype 5 ^23^: AAV5.hSyn.HI.eGFP-Cre.WPRE.SV40 (Addgene; #105540). Sparse labelling was obtained using Cre-inducible, double inverted open (DIO)-reading frame AAV2-hSyn-DIO-mCherry (Addgene; #50459) diluted to 1.5×10^11^ viral particle/mL for neuronal morphology ^24^.

For Nr4a1 overexpression virus, Nr4a1 sequences were obtained from Origene (#MR209316), PCR amplified (Phusion DNA polymerase; New England Biolabs) and cloned into the NheI and NcoI restriction site of the AAV-EF1a-DIO-EYFP vector. The AAV-DIO-Nr4a1-EYFP was packaged into AAV (serotype 9) at UMB Virus Vector Facility. For verification of Nr4a1 overexpression, 300-400 ng of cDNA from VP was synthesized using the reverse transcriptase iScript cDNA synthesis kit (Bio-Rad). mRNA expression changes were measured using qRT-PCR with PerfeCTa SYBR Green FastMix (Quanta). Quantification of mRNA changes was performed using the − ΔΔCT method described previously ^17^.

Due to the large size of SpCas9 (4.5 kb), which limits its packaging into an AAV, we chose the SaCas9 CRISPR protein (3.1 kb). By facilitating its packaging into an AAV, its *in-vivo* delivery for genome editing is thus facilitated ^25^. Additionally, studies showed that SaCas9 possesses higher cleaving activity over SpCas9 ^26^. Moreover, SaCas9 recognizes a 5′-NNGRRT-3′ PAM sequence which is more specific than the SpCas9 PAM sequence. The AAV-CMV-DIO-SaCas9 was packaged into AAV (serotype 9) at UMB Virus Vector Facility (**Supplementary Figure 4a, b**; see **Supplementary Methods** for detailed procedure).

We further verified CRISPR knockdown efficacy using qRT-PCR: 300-400 ng of cDNA from VP was synthesized as described above and mRNA expression was quantified using the − ΔΔCT method.

### RNA sequencing and bioinformatics

Bulk ventral pallidum RNA was extracted with the RNeasy Micro kit (Qiagen: 74004; see below for detailed extraction procedure). For RNA sequencing, only samples with RNA integrity numbers >8 were used. Samples were submitted in biological quadruplicates for RNA sequencing at the UMSOM Institute for Genome Sciences (IGS) and processed as in ^14^.

Libraries were prepared from 90-ng of RNA from each sample using the NEBNext Ultra kit (New England BioLabs). Samples were sequenced on an Illumina HiSeq 4000 with a 75 bp paired-end read. An average of 86 million reads were obtained for each sample. Reads were aligned to the mouse genome (Mus_musculus. GRCm38) using TopHat2 ^27^ (version 2.0.8; maximum number of mismatches= 2; segment length= 30; maximum multi-hits per read= 25; maximum intron length= 50,000). The number of reads that aligned to the predicted coding regions was determined using HTSeq ^28^. Significant differential expression was assessed using DEseq. RNA-sequencing data are available through the Gene Expression Omnibus database (GEO accession number: GSE199725), the gEAR portal ^29^ (https://umgear.org/p?s=27207e4b) as well as in **Supplementary Table 1**.

Gene ontology (GO) functional enrichment analysis was performed on significantly differentially expressed (SDE) genes with Cytoscape software (v.3.6.1) ^30^ using the BiNGO plugin ^31^. Follow up transcriptional regulator network were obtained with iRegulon ^32^ by analyzing genes from significant “Cellular Component” GO terms.

### RNA extraction and qRT-PCR

RNA extraction and quantitative RT-PCR was performed as previously described ^33^. Briefly, tissue punches were collected using a 14-gauge punch targeting the VP (under the anterior commissure (post limb) at approximately AP +0.2 mm from bregma), 24-h after the last self-administration session or 3 weeks after viral infusion and stored at -80°C. RNA was extracted using Trizol (Invitrogen) and the E.Z.N.A MicroElute Total RNA Kit (R6831-01, Omega) with a DNase step (#79254, Qiagen). RNA quantity was measured using a Nanodrop (Thermo Scientific). Three to four hundred nanograms of complementary DNA was synthesized using a reverse transcriptase complementary DNA synthesis kit (iScript, Bio-Rad). Changes in mRNA expression were measured using quantitative PCR (Biorad) with PerfeCTa SYBR Green FastMix (#95072, Quantabio). Quantification of mRNA changes was performed using the 2–ΔΔCt method, using glyceraldehyde 3-phosphate dehydrogenase (Gapdh) as a housekeeping gene.

The list of primers used is available in **Supplementary Table 3**. From the qRT-PCR validation experiment, one male-cocaine and one female-cocaine samples were excluded due to low RNA concentrations.

### Projection-specific mRNA expression

Polyribosome immunoprecipitation was achieved as described in ^14, 33^. Briefly, pooled tissue from RiboTag (RT)^+/+^ mice with virally mediated Cre expression in VP→MDT neurons (n= 4-5 mice per sample) was homogenized and 800-μL of the supernatant was incubated in HA-coupled magnetic beads (Invitrogen: 100.03D; Covance: MMS-101R) overnight at 4°C. Magnetic beads were then washed in high salt buffer. Following TRK lysis buffer addition, RNA was extracted with the RNeasy Micro kit (Qiagen: 74004). For input, 50 ng of RNA was used and 1-2 ng of RNA from immunoprecipitated samples were amplified using the Low RNA Input kit (NanoString Technologies®). All samples were then processed with the nCounter Master Kit (NanoString Technologies®) by UMSOM IGS on a custom-made gene expression Code set (see **Supplementary Table 3** for primer sequences). Data were analyzed with nSolver Analysis software ^14^.

### In-situ *hybridization*

Coronal brain slices (16-18µ m) were prepared using a cryostat (Leica) and fluorescent *in-situ* hybridization was performed using the RNAscope® kit following manufacturer’s instructions (Acdbio, Advanced Cell Diagnosis). Briefly, slide-mounted sections were post-fixed in 10% formalin and dehydrated in ethanol. RNA probes targeting Cre (#312281) and Nr4a1 (#423341- C3) were applied and DAPI staining was used to identify cell nuclei.

### Immunohistochemistry

Mice were perfused with 0.1M PBS and 4% paraformaldehyde and brains were post-fixed for 24 h. For dendritic spine morphology and virus placement, 100 μm sections were blocked in 3% normal donkey serum and 0.3% Triton X-100 in PBS for 30 min, then incubated in chicken anti-mCherry (1:500; Novus Biologicals, NBP2 25158) and in rabbit anti-GFP (1:500; Cell Signaling #2555) overnight at 4°C as in ^34, 35^. For virus validation a mouse monoclonal anti-Nr4a1 antibody (1:1000; #sc-365113, Santa Cruz) or a polyclonal rabbit anti-SaCas9 (1:1000; #ab203933, Abcam) were used. Following eight 1 h-PBS washes slices were incubated overnight in anti-chicken-Cy3 (1:1000, Jackson Immuno; #703-165-155) and anti-rabbit-Alexa 488 (1:1000, Jackson Immuno; #111-545-003) or anti-mouse Cy2 (1:1000, Jackson Immuno; #715-225-150), anti-rabbit Cy5 (1:1000, Jackson Immuno; #711-175-152) depending on the condition at 4°C. Finally, following eight 1 h-PBS washes, slices were mounted with Aqua-Poly/Mount (Polysciences) mounting media.

### Imaging

RNA puncta counting: Olympus FV500 Confocal Microscope was used to image immunofluorescence. VP images were taken at 40x magnification from 4 hemispheres per animal out of 3 sections between AP: +0.3 and +0.0 from bregma (approx.). Each VP image sample contained 6 stacked images with 1.5µ m step. Counting was performed using Image J software (Fiji plugin) ^36^ by merging all 6 stacked images. Neurons from each projection were identified by the presence of Cre puncta within the boundaries of the VP as described by ^37^. For colocalization, 300-400 Cre-positive cells were assessed per animal and the proportion of Cre + Nr4a1 double labeling was calculated. For puncta per cell measurements, individual puncta were counted from 120 Cre + Nr4a1 positive cells per mouse. Counting was performed by experimenters blind to the treatment/group conditions.

Dendritic spine counting: Spine images (from VP→MDT neurons only) were acquired on a laser-scanning confocal microscope (Leica SP8) with 0.2-μm increments Z-stacks using a 63x objective with 2x digital magnification ^14, 34^ and quantified with Neuron Studio software (Mt Sinai School of Medicine) ^38^. Secondary dendrites were selected for imaging and subsequent counting. For all morphological analyses 2–5 cells, taken between AP: +0.3 and +0.0 from bregma (approx.) within VP boundaries described by ^37^, were averaged per animal. The expression of Cre-dependent mCherry insured circuit specificity. Morphology was blindly assessed.

### Statistical analysis

Graphpad Prism 8.4.3. software was used for statistical analysis. Normality was assessed with Bartlett’s test. 2-way RM-ANOVAs and 3-way RM-ANOVAs (mixed-effects model) were run with Sidak, and Tukey posthoc tests respectively. Unpaired two-tailed t-tests were used when appropriate. Samples were excluded if animals did not acquire cocaine self-administration, if not detected (molecular analysis), had inappropriate viral placement (i.e., no/unilateral expression, off-target Cre injection), or failed Grubbs’ outlier test (See **Supplementary Methods** for detailed numbers). No more than 3 animals per group were excluded. Sample sizes were determined from previous studies using self-administration in mice, RNA sequencing, cell-type-specific RNA isolation and neuronal morphology ^14, 15, 24^. In figure legends: *p≤ 0.05, **p< 0.01, ***p< 0.001. All graphs represent mean ± s.e.m. Individual values are plotted to report that variation and variance is similar between groups that are compared. Detailed statistics are available in **Supplementary Material 1** and in figure captions of supplementary figures for supplementary data statistics. Data will be made available by the authors upon reasonable request. RNA sequencing data are available through the Gene Expression Omnibus database (GEO accession number: GSE199725).

## Results

### Transcriptome profiling of ventral pallidum reveals cocaine-induced changes in structural plasticity-related genes

To characterize the molecular alterations associated with cocaine intake, we trained male mice to self-administer cocaine (or saline) for 10 days. Mice showed significantly higher numbers of cocaine infusions compared to saline starting at day 3: 2-way RM-ANOVA: Day x Drug: F_(9, 54)_= 2.7; p= 0.011; Sidak posthoc: p< 0.01 at least (**Figure 1a**). VP tissue was collected 24-hr after the last drug intake session and processed for RNA sequencing. Out of the 32,787 genes identified with RNA sequencing, 363 genes (1.11%) were differentially expressed in mice taking cocaine compared to saline: FDR< 0.05 (**Supplementary Table 1** for detailed RNA sequencing results). From this subset of genes, 170 (46.8%) were downregulated and 193 (53.2%) were upregulated (**Figure 1b**). Gene ontology (GO) has been conducted using the 363 significantly differentially expressed (SDE) genes. At the “Biological process” level, several GO terms were associated with neuronal morphology and neuronal communication. Among them we found, ‘Signal transduction’, ‘Cell communication’, ‘Synaptic transmission’, ‘Cell projection organization’, ‘Cytoskeleton organization’, ‘Cell morphogenesis’, ‘Regulation of neuron projection development’, ‘Neuron projection morphogenesis’, ‘Actin cytoskeleton organization’, ‘Regulation of synaptic plasticity’ (**Figure 1c;** see **Supplementary Figure 1a** for details).

**Figure 1.**
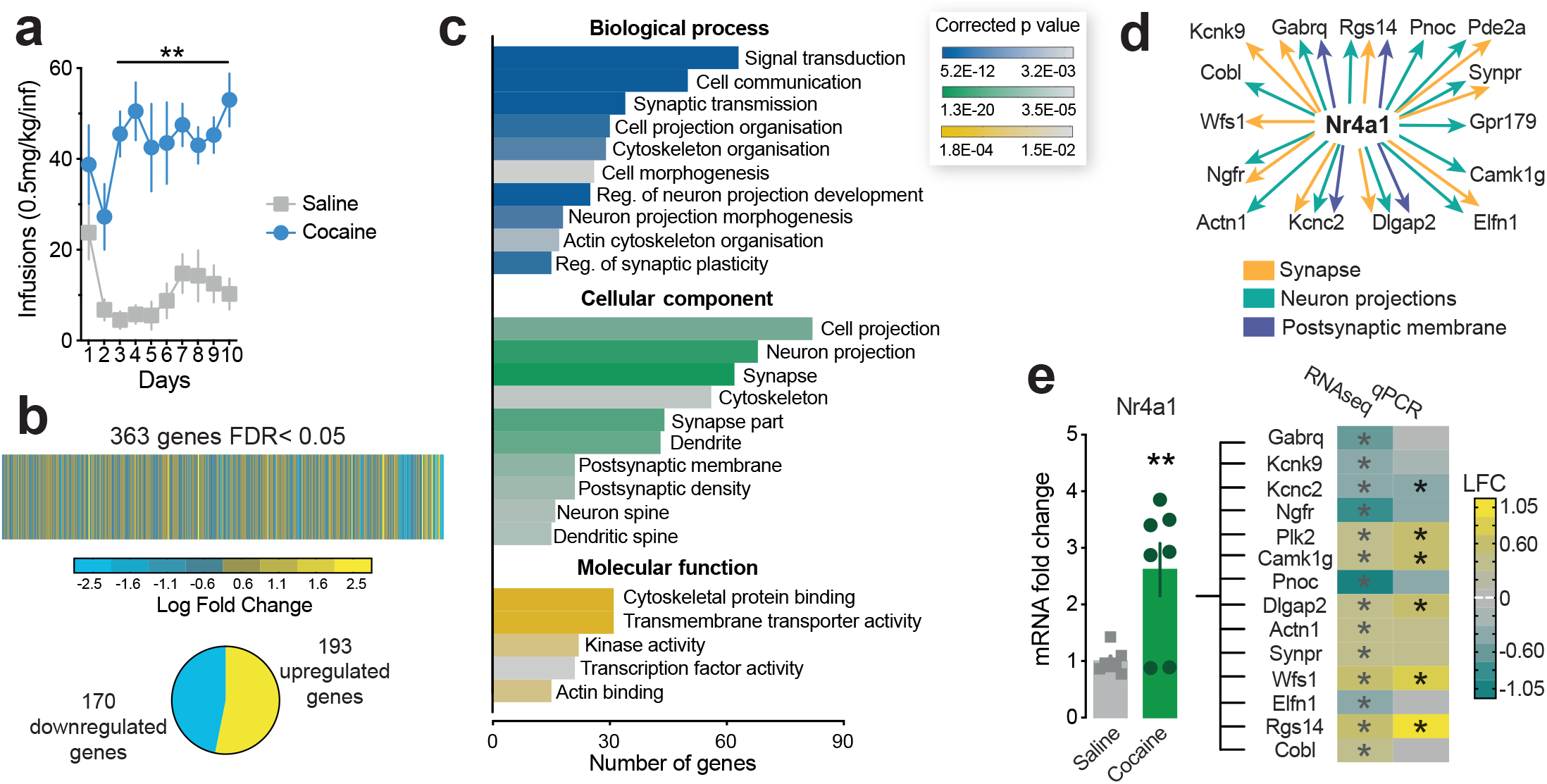
Cocaine intake alters structural plasticity-related molecules in the ventral pallidum. **a**: Number of infusions acquired during the 10 days of self-administration (cocaine vs. saline: p< 0.01 from day 3 onward). n= 4 male mice/group; **b**: Ventral pallidum tissue was collected 24 hrs following self-administration and used for RNA sequencing. Heat map of the 363 genes showing significant False Discovery Rate (FDR; p< 0.05); **c**: Gene Ontology analysis (GO) revealed a high number of significant GO terms related to neuronal morphology and structural plasticity. Of note, neuronal morphology-related GO terms represented 40.6% of the “Cellular Component” GO terms. These morphology-related GO terms included 200 unique genes (i.e., 55% of all the significantly differentially expressed (SDE) genes); **d**: Transcriptional regulator network analysis showed Nr4a1 (Nurr 77) is a common regulator for SDE genes from Synapse, Neuron projections, Postsynaptic membrane GO terms; **e**: Left bar graph: qRT-PCR analysis confirmed RNAseq data and showed a significant increase of *Nr4a1* expression in the VP of male mice 24 hrs following 10 days of cocaine self-administration (p< 0.01). Right heat maps: qRT-PCR analyses further confirmed significant changes in multiple Nr4a1 target genes initially identified with RNAseq and bioinformatics analyses (p< 0.05 at least). For RNAseq n= 4/group and for qRT-PCR, n= 7/group. Detailed statistics are available in **Supplementary Table 2**. See also **Supplementary Figure 1**.

At the “Cellular component” level, neuronal morphology-related GO terms represented 40.6% of the “Cellular Component” GO terms. There were 200 unique genes in the morphology-related GO terms, which represents 55% of all the SDE genes and 63% of the SDE genes used for the GO analysis at the Cellular component level. Among the significant neuronal morphology-related GO terms, we found: ‘Cell projection’, ‘Neuron projection’, ‘Synapse’, ‘Cytoskeleton’, ‘Synapse part’, ‘Dendrite’, ‘Postsynaptic membrane’, ‘Postsynaptic density’, ‘Neuron spine’, ‘Dendritic spine’ (**Figure 1c**; see **Supplementary Figure 1a** for details). Together, our findings related to structural plasticity and dendritic spines are consistent with described drug-induced changes in neuronal activity ^1, 3, 12^.

Finally, at the “Molecular function” level, among the relevant significant GO terms, we found: ‘Cytoskeletal protein binding’, ‘Transmembrane transporter activity’, ‘Kinase activity’, ‘Transcription factor activity’, ‘Actin binding’ (**Figure 1c**; see **Supplementary Figure 1a** for details).

### Nr4a1 as a potential regulator of structural plasticity

Our GO term analysis suggests important alterations in structural plasticity-related functions. To find potential transcriptional mechanisms underlying these changes, we used the transcriptional regulator analysis iRegulon ^32^ on genes from significant GO terms at the “Cellular component” level. Among the regulators highlighted by our analysis, direct motif similarity was found between the transcription factor *Nr4a1* (Nuclear receptor subfamily 4, group A, member 1, or *Nur77*) and genes implicated in various cellular functions: 13 genes (19.1%) from the GO term ‘Neuron projection’ (Normalized Enrichment Score (NES): 3.165), 10 genes (16.1%) from the GO term ‘Synapse’ (NES: 3.455) and 4 genes (19%) from the GO term ‘Postsynaptic membrane’ (NES: 3.172) (**Figure 1d**). These 3 GO terms associated with *Nr4a1* appeared particularly relevant as they regrouped a large number of genes (84 unique genes; 23% of our SDE genes) and the terms ‘Synapse’ and ‘Neuron projection’ presented the lowest *p* values at the “Cellular component” level. (**Supplementary Figure 1a**). Moreover, Nr4a1 is an orphan nuclear receptor that acts as a transcription factor, known to be induced by psychostimulants and capable of activity-dependently reshape synaptic functions ^39-41^. Finally, this transcription factor was significantly upregulated in our RNA sequencing analysis (**Supplementary Table 1**). Together, these different factors supported *Nr4a1* as a suitable target for our subsequent investigations. Since our analyses pointed toward changes in *Nr4a1* and its target genes in neuronal morphology adaptations, we evaluated these changes in a new cohort of male mice using qRT-PCR (**Figure 1e; Supplementary Figure 1b; Supplementary Table 2** for primer sequences). Quantitative RT-PCR of VP tissue confirmed a significant increase of *Nr4a1* mRNA expression following cocaine intake (+162%; **Figure 1e**). Moreover, 42.9% of *Nr4a1* gene-targets identified by our transcriptional regulator analysis and well-characterized findings from the literature^41^ were significantly changed following cocaine intake (**Figure 1e; Supplementary Table 2**). As our transcriptomic profiling was performed in male mice, we ran the same qRT-PCR validation experiment in a cohort of female mice. We did not observe significant changes in Nr4a1 expression or any of its target genes suggesting *Nr4a1*-related changes were specific to males (**Supplementary Figure 1c, d**). Thus, all the subsequent experiments described below were done in male mice exclusively.

### Ventral pallidum to mediodorsal thalamus neurons molecular profiling

In order to investigate whether changes in Nr4a1 expression occurred in all VP neurons or were restricted to specific neuronal subpopulations, we focused on three main VP projection populations that are important for reward processing: the ventral tegmental area (VTA), the lateral habenula (LHb) and the mediodorsal thalamus (MDT) (**Figure 2a**). To label specific subpopulations, we took advantage of the well described retrograde properties of AAV5 vectors (AAV5-Cre-GFP) ^23, 42^. We infused each output structure in distinct groups of male mice before submitting them to our self-administration procedure (**Supplementary Figure 2a**). Twenty-four hours later VP tissue was collected for *in situ* hybridization. When looking at the VP neurons projecting to the VTA, the proportion of cells expressing Nr4a1 (Cre + Nr4a1) was significantly decreased in mice taking cocaine (29.8%) compared to saline controls (42.9%): Chi^2^= 6.895, p= 0.0086. This proportion was not changed in VP neurons projecting to the LHb between animals taking cocaine (46.7%) or saline (44.8%): Chi^2^= 0.162, p= 0.69. Interestingly, the proportion of VP neurons projecting to the MDT expressing *Nr4a1* was increased in mice taking cocaine (63.2%) compared to saline controls (45%): Chi^2^= 13.091, p= 0.0003 (**Figure 2b**). Evaluating the number of *Nr4a1* puncta per cell in these 3 projection neurons revealed that only the VP neurons projecting to the MDT showed increased (+76%) Nr4a1 expression levels following cocaine self-administration: 2-way ANOVA: Projection x Drug: F_(2, 24)_= 3.54; p= 0.045; Sidak posthoc: p< 0.05 (**Figure 2c**). Together, our circuit analysis showed that only VP→MDT neurons, had increases for both *Nr4a1* expression levels and the proportion of cells expressing *Nr4a1* after cocaine intake. We thus chose this specific projection population for further investigation.

**Figure 2.**
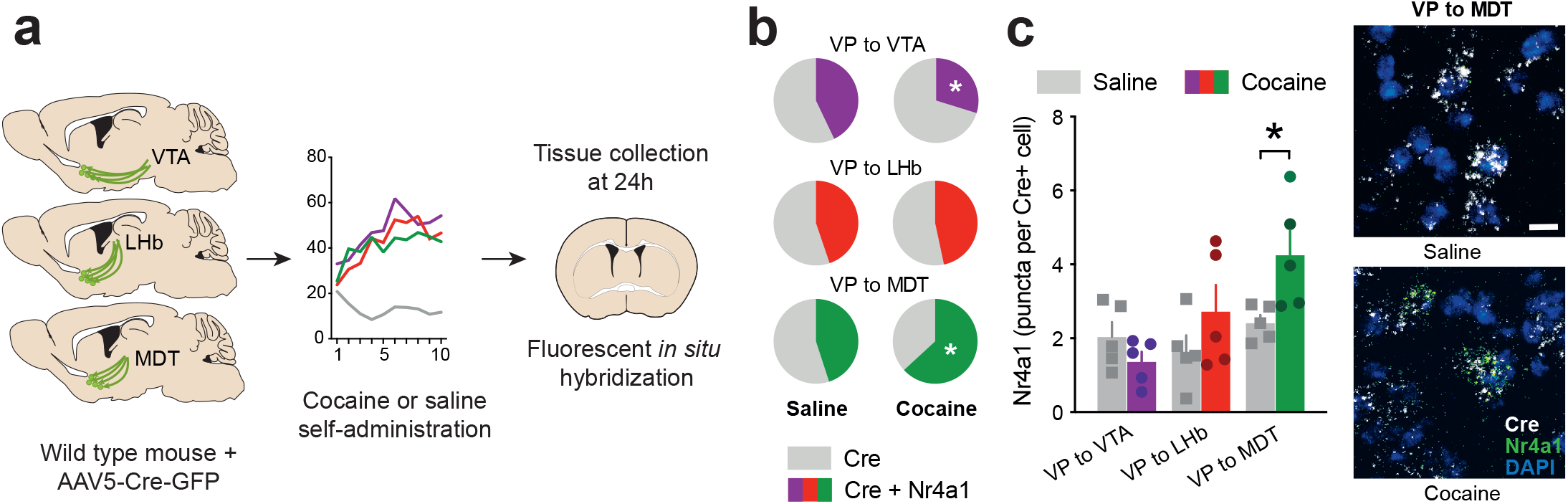
Cocaine intake increases *Nr4a1* expression specifically in VP neurons projecting to the MDT. **a**: Schematic of the experimental design: 3 different groups of mice received an infusion of retrograde (AAV5) Cre-GFP vector in either the ventral tegmental area (VTA), the lateral habenula (LHb) or the mediodorsal thalamus (MDT). Male mice were then subjected to either cocaine (0.5 mg/kg/inf.) or saline self-administration for 10 days. Twenty-four hours later, brain sections containing the VP were collected for fluorescent *in situ* hybridization; **b**: Proportion of Cre-positive cells expressing Nr4a1 among all the Cre-positive cells in each group. The proportion of cells expressing Nr4a1 was significantly lower in VP cells projecting to the VTA (p< 0.01), unchanged in cells projecting to the LHb and significantly higher in cells projecting to the MDT (p< 0.001); **c**: Nr4a1 puncta density (puncta per Cre-positive cell) was assessed for each group. Only VP cells projecting to the MDT showed a significantly higher Nr4a1 expression level following cocaine intake compared to saline control (p< 0.05). n= 5/group; density was measured from 120 Cre-positive cells/animal. Right panel: representative image from VP cells projecting to the MDT from each group, scale bar= 20 µ m. See also **Supplementary Figure 2**.

As Nr4a1 is involved in structural plasticity and dendritic spine remodeling ^41^, we chose to pursue our studies with a 4-hour time-point that would allow us to assess dendritic spine dynamics ^43^. In a cohort of male RiboTag mice subjected to cocaine self-administration, VP→MDT neurons were immunoprecipitated and mRNA levels were measured with a multiplexed assay (**Figure 3a, Supplementary Figure 2b**). We selected 52 genes to both characterize the identity of VP→MDT neurons and measure changes in Nr4a1 target genes and known structural plasticity-related molecules (**Supplementary Table 2** for primer sequences).

**Figure 3.**
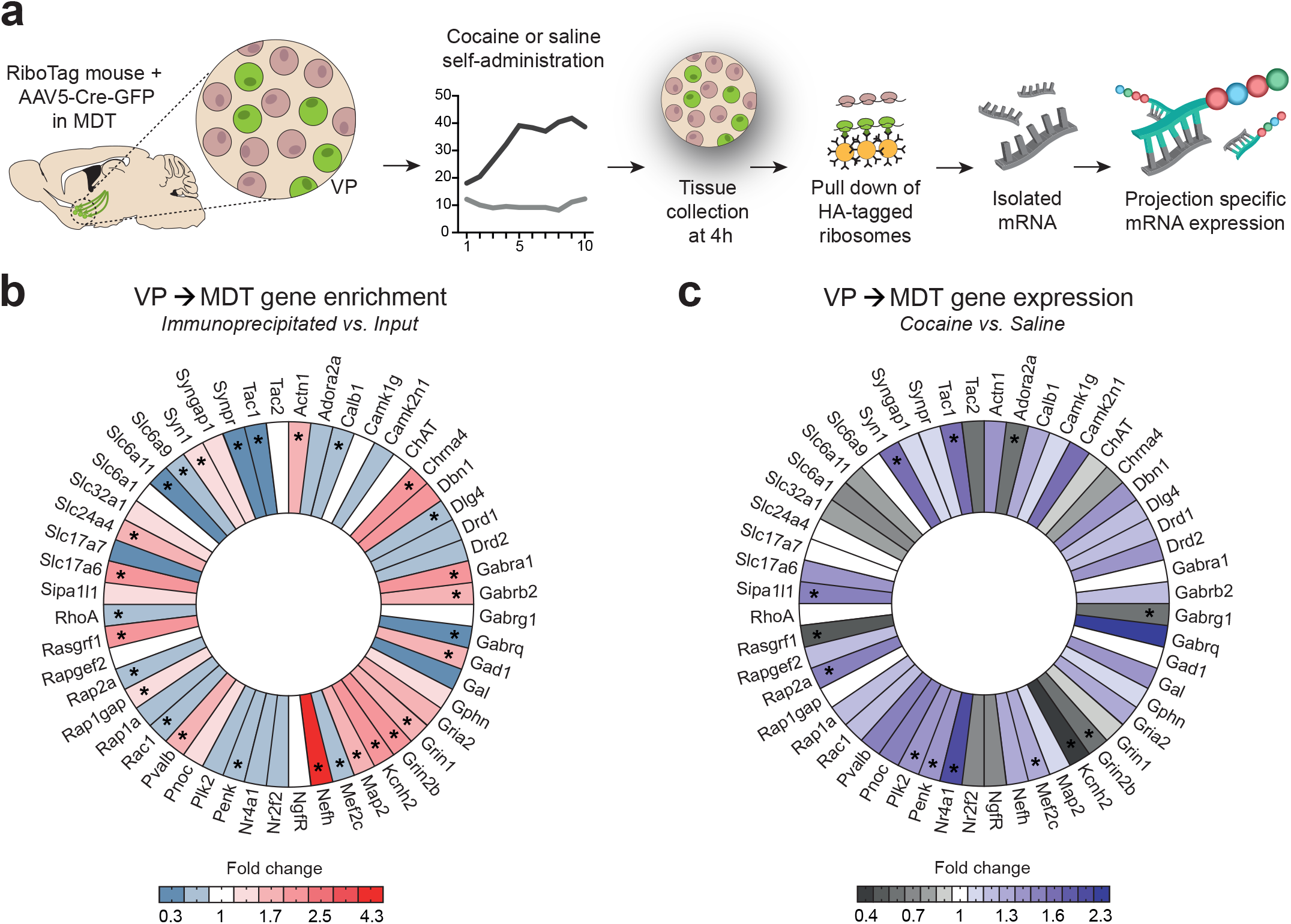
Transcriptional characterization of VP neurons projecting to the MDT in controls and following cocaine intake. **a**: Schematic of the experimental design: RiboTag male mice were injected with retrograde (AAV5) Cre-GFP vectors. Animals were subjected to 10 days of cocaine (0.5 mg/kg/inf.) or saline self-administration and VP tissue was collected 4 hrs following the last session. Tagged polyribosomes from VP to MDT neurons were immunoprecipitated, mRNA was isolated and gene expression was analyzed using a multiplexed gene expression assay. n= 4-5 mice/sample, 6 samples/condition; **b**: Heat map representing gene enrichment in VP to MDT neurons in saline control mice (p≤ 0.05 at least); **c**: Heat map showing differences in gene expression, specifically in VP to MDT neurons, in mice taking cocaine compared to mice taking saline (p≤ 0.05 at least). Detailed statistics are available in **Supplementary Table 3**. See also **Supplementary Figure 2**.

Assessment of gene enrichment in the VP→MDT neurons from saline mice (**Figure 3b; Supplementary Table 3**) indicated that this cell-population expressed higher parvalbumin (*Pvalb*) and lower calbindin (*Calb1*) levels than the general VP. The enrichment of *Gad1* (Glutamate Decarboxylase 1) and *Slc17a6* (*Vglut2*; Vesicular Glutamate Transporter 2) indicated that this projection contained both GABAergic and glutamatergic neurons ^1, 12^. *Chrna4* (Cholinergic Receptor Nicotinic α 4 Subunit) was highly enriched in these cells, but levels of *ChAT* (Choline Acetyltransferase) were comparable to bulk VP. VP→MDT cells also expressed higher levels of N-methyl-D-aspartate receptor subunits 1 and 2b (*Grin1* and *Grin2b*) as well as GABAergic receptor subunits α 1 and β 2 (*Gabra1* and *Gabrb2*). Additionally, VP→MDT neurons were highly enriched in Neurofilament Heavy Chain (*Nefh*; +332%). Finally, enrichment for *Actn1* (Actinin α 1), *Syn1* (Synapsin I), *Map2* (Microtubule Associated Protein 2), *Rap1gap* (RAP1 GTPase Activating Protein), *Rasgfr1* (Protein Phosphatase 1 Regulatory Subunit 9B), *Slc24a4* (Sodium/Potassium/Calcium Exchanger 4) and *Kcnh2* (Potassium Voltage-Gated Channel Subfamily H Member 2 were significantly increased in VP→MDT neurons (**Supplementary Table 3** for detailed statistics). Oppositely, these cells had reduced proenkephalin (*Penk*) and substance P (*Tac1*) levels suggesting low peptidergic transmission. Enrichment in *Dlg4* (Discs Large MAGUK Scaffold Protein 4, or PSD-95), *Gabrq* (GABA_A_ Receptor Subunit Theta), *Rac1* (Rac Family Small GTPase 1), *Rap2a* (Ras-Related Protein Rap-2a), *RhoA* (Ras Homolog Family Member A), *Synpr* (Synaptoporin), Mef2c (Myocyte Enhancer Factor 2C), *Slc6a9* (Sodium- And Chloride-Dependent Glycine Transporter 1) and *Slc6a11* (Sodium- And Chloride-Dependent GABA Transporter 3) was significantly decreased in VP→MDT neurons (**Supplementary Table 3** for detailed statistics) (**Figure 3b**). Of note, these enrichment patterns were largely identical in mice taking cocaine, confirming that these genes were stable markers of this neuronal population (**Supplementary Figure 2c**).

We then compared changes in gene expression due to cocaine intake specifically in VP→MDT neurons (**Figure 3c**). We first confirmed increased *Nr4a1* levels after cocaine intake in this subpopulation at the 4h time-point (as well as in the whole VP: **Supplementary Figure 2c**). The *Nr4a1* upstream regulator *Mef2c* was also increased along with the *Nr4a1* downstream target *Plk2* (Polo-Like Kinase 2). *Sipa1l1* (Signal Induced Proliferation Associated 1 Like 1; or SPAR), a downstream target of Plk2, showed high mRNA levels associated with high *Rap2a* levels and low mRNA levels of *Rasgrf1*. Additionally, *Syn1* levels were increased, supporting changes in synaptogenesis and neurotransmitter release. Although basal enrichment of *Penk* and *Tac1* in VP→MDT neurons was low, cocaine intake significantly increased levels of both these molecules (**Supplementary Table 3** for detailed statistics). Finally, enrichment in *Adora2a* (Adenosine A2a Receptor), *Gabrg1* (GABA_A_ Receptor Subunit Gamma-1), Grin2b and Kcnh2 were significantly decreased in this neuronal subpopulation after cocaine intake (**Supplementary Table 3** for detailed statistics).

### Pallido-thalamic Nr4a1 expression bidirectionally regulates cocaine-related behavior and dendritic spine density

Knowing that Nr4a1 is involved in structural plasticity and that its expression was increased in VP→MDT neurons following cocaine intake, we interrogated how artificially increasing *Nr4a1* levels (**Supplementary Figure 3a, b**) in this specific cell-population would impact cocaine intake and dendritic spine dynamics (**Figure 4a, f** and **Supplementary Figure 3c**). First, male mice receiving Nr4a1 overexpression in VP→MDT neurons showed no difference in natural reward (water) self-administration compared to eYFP controls: 2-way RM-ANOVA: Session x Virus: F_(7, 189)_= 0.799; p= 0.589 (**Supplementary Figure 3d**). Cocaine and saline self-administration were then compared in mice with Nr4a1 or eYFP overexpression in VP→MDT neurons. A 3-way ANOVA showed a significant Day x Drug interaction: Day x Drug: F_(9, 41)_= 8.803; p< 0.0001 but no Day x Virus interaction: Day x Virus: F_(9, 171)_= 0.468; p= 0.894, suggesting that both mice with eYFP and Nr4a1 overexpression were taking more doses of cocaine compared to their saline counterparts, although there were no differences in acquisition rate between the two groups taking cocaine. Tukey’s multiple comparison showed that eYFP mice were taking more doses compared to their respective control group from day 4 onward (p< 0.05 at least). This difference was observed in mice with Nr4a1 overexpression at day 5 and day 8 onward (p< 0.05) (**Figure 4b**). Similar results were obtained when assessing active and inactive responses, further indicating that both groups acquired and discriminated the active and inactive ports similarly (**Supplementary Figure 3e**). Mice were then tested for cocaine seeking behavior. Both mice with Nr4a1 or eYFP overexpression in VP→MDT neurons showed significant seeking behavior compared to their respective saline controls. However, mice with Nr4a1 overexpression previously taking cocaine showed higher levels of active responses compared to mice with eYFP overexpression previously taking cocaine: 2-way ANOVA: Drug: F_(1, 25)_= 37.23; p< 0.0001; Drug x Virus: F_(1, 25)_= 3.93; p= 0.058; Tukey posthoc: p< 0.05 at least (**Figure 4c**). Following seeking test, mice were evaluated for extinction in similar test conditions. A 3-way ANOVA showed a significant Session x Drug interaction: Session x Drug: F_(4, 11)_= 9.005; p< 0.01 but no Session x Virus interaction: Session x Virus: F_(4, 76)_= 0.024; p= 0.998 suggesting again that both mice with *eYFP* and *Nr4a1* overexpression previously taking cocaine were making more active responses compared to their saline counterparts, although there were no differences in extinction rate between the two groups previously taking cocaine. Tukey’s multiple comparison showed that both mice with *eYFP* and *Nr4a1* overexpression, previously taking cocaine, made more active responses than their respective saline controls on Session 1: p< 0.01 (**Figure 4d**). Finally on the next day, mice were tested for drug induced reinstatement. A 2-way ANOVA highlighted a Drug effect suggesting that a low dose of cocaine increased the number of active responses in mice previously taking cocaine: 2-way ANOVA: Drug: F_(1, 25)_= 11.06; p= 0.002. A Tukey multiple comparison showed no significant difference between *eYFP* and *Nr4a1* mice exposed to cocaine (p=0.555) but revealed however that only mice previously exposed to cocaine with Nr4a1 overexpression showed significantly increased responding compared to their respective controls (p= 0.050). This was not observed for similar conditions in mice with eYFP overexpression (p= 0.245) (**Figure 4e**). Four hours following the drug-induced reinstatement test, mice were perfused and dendritic spine density was assessed in VP→MDT neurons specifically (**Figure 4g**). A 2-way ANOVA showed a significant Drug x Virus interaction: Drug x Virus: F_(1, 25)_= 5.841; p= 0.023. Tukey posthoc test revealed that spine density was higher in mice with eYFP overexpression previously taking cocaine compared to their respective saline controls (+167.2%; p< 0.01) but also compared to mice with Nr4a1 overexpression previously taking cocaine (+176.5%; p< 0.01) (**Figure 4h**). Spine subtype analysis revealed that overall, these changes mostly affected thin spines. (**Supplementary Figure 3f**).

**Figure 4.**
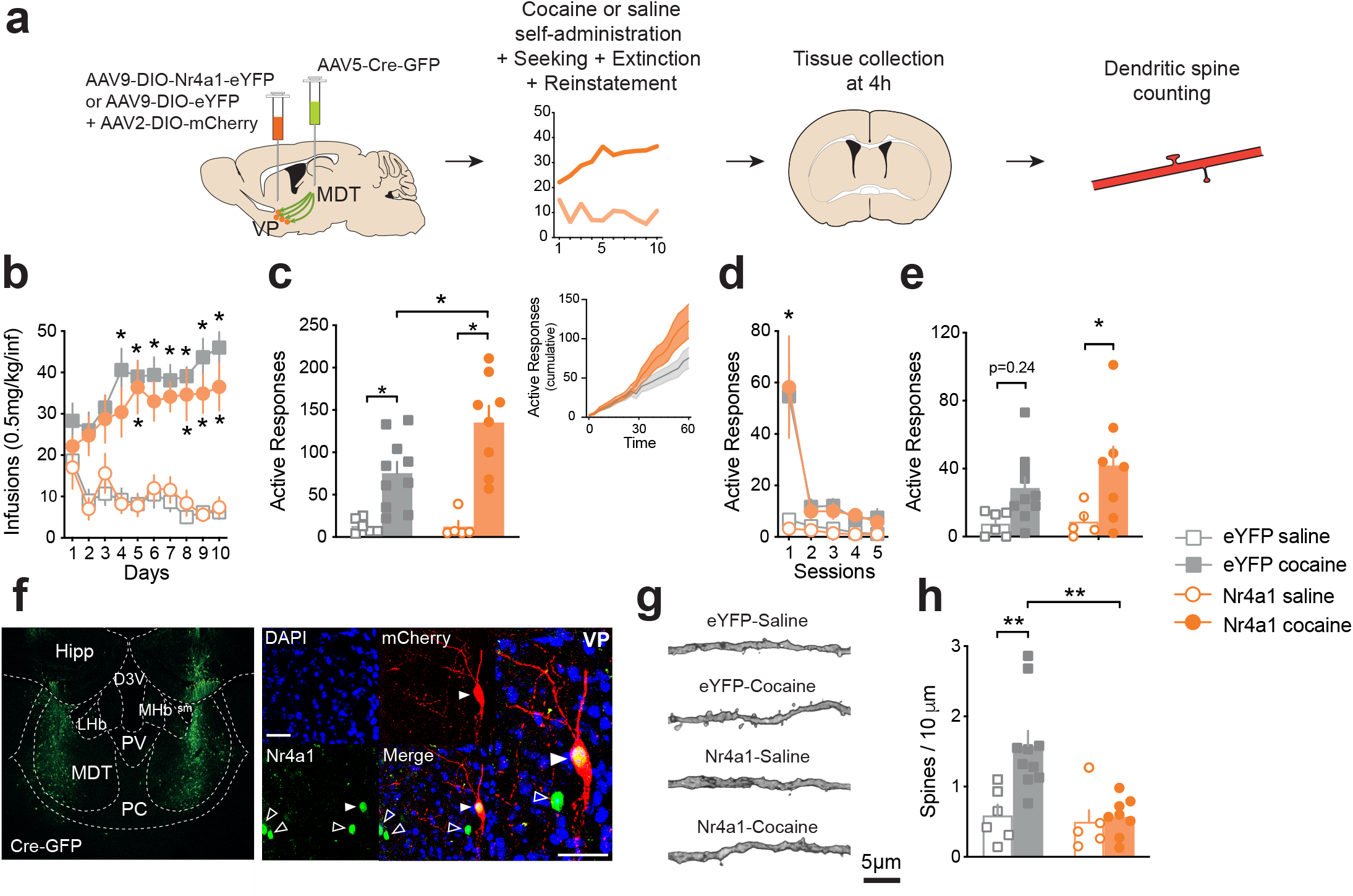
Overexpression of Nr4a1 in VP to MDT neurons enhanced relapse-like behaviors and reduced dendritic spine density. **a**: Schematic of the experimental design: Male mice received an infusion of retrograde (AAV5) Cre-GFP vector in the MDT and an infusion of DIO-Nr4a1-eYFP (or DIO- eYFP) + DIO-mCherry in the VP prior to 10 days of cocaine (0.5 mg/kg/inf.) or saline self- administration. On day 11, mice were subjected to seeking test (under extinction condition) as well as 5 additional extinction sessions. On day 12, mice were tested for drug-induced reinstatement. Four hours following this last test, tissue was collected for spine density evaluation. n= 6 eYFP-saline, 10 eYFP-cocaine, 5 Nr4a1-saline, 8 Nr4a1-cocaine; **b**: Both groups show significantly higher intake compared to their respective controls (p< 0.05 at least); **c**: Both groups showed significant seeking behavior compared to their respective saline controls. However, mice with Nr4a1 overexpression showed higher levels of active responses compared to eYFP overexpression mice (p< 0.05 at least). Insert represents cumulative active response in cocaine mice during the 1 hr seeking test (5 min bins); **d**: Active responses during extinction test. Both groups show similar rate of extinction (p< 0.05 from respective saline controls); **e**: Animals previously taking cocaine were exposed to non-contingent cocaine for drug-induced reinstatement (7.5 mg/kg, i.p.; saline controls received saline, i.p.). While eYFP-cocaine mice showed a trend in increased active responses, Nr4a1-cocaine mice made significantly more active responses compared to their respective control (p< 0.05); **f**: Left panel: Representative image of MDT injection site for retrograde (AAV5) Cre-GFP viral vector. Right panel: Representative image of Nr4a1 positive cells in mice with Nr4a1 overexpression and sparse DIO-mCherry labelling in the VP. Closed arrowhead: Nr4a1+mCherry positive cell, open arrowhead: Nr4a1 positive cell. Scale bars= 25 µ m; **g**: Representative image of dendritic spines from sparsely labelled VP to MDT neurons from each group. Scale bar= 5 µ m; **h**: Spine density was significantly increased in control mice taking cocaine while Nr4a1 overexpression blocked this increase (p< 0.01). See also **Supplementary Figure 3**.

Together, this suggested cocaine intake induced *Nr4a1* expression and increased thin spine density. However, we found *Nr4a1* overexpression decreased spinogenesis. To provide further insight into these apparent contradictions, we explored a mechanism likely to explain our observations. Bioinformatics analyses and subsequent qPCR validation at the VP level highlighted significant changes in several *Nr4a1* gene targets such as *Plk2, Dlgap2* (DLG Associated Protein 2) and *Rgs14* (Regulator Of G-Protein Signaling 14). These genes are involved in spine remodeling through the Ras/Raf GTPase signaling pathway ^44-46^. We thus assessed at the VP level, the potential involvement of the small GTPase RAP2 (Ras-Related Protein) for its role in inhibiting dendritic spine development ^47^. We exposed mice to cocaine for 10 days (20mg/kg, ip.) and collected VP tissue 24h later. The RAP2 activity assay revealed a significant increase in RAP2 activity in the VP: t_12_ = 2.272; p= 0.0423 (**Supplementary Figure 3g**). This suggests that cocaine self-administration induces spinogenesis inhibition mechanisms in control conditions, that are likely counterbalanced by compensatory mechanisms allowing for optimal spine regulation ^44^. Because our viral manipulation induced prolonged and supraphysiological expression of *Nr4a1* in VP→MDT neurons, we likely disrupted this balance in spine regulation mechanisms to produce blunted spinogenesis.

Our *in-situ* hybridization experiment showed a reduced proportion of cells expressing *Nr4a1* in VP→VTA neurons (**Figure 2b**). We performed a similar overexpression study in this cell population to assess whether preventing reduction in *Nr4a1* expression levels would affect cocaine-related behaviors. However, we found that *Nr4a1* overexpression in this circuit had no effect on self-administration acquisition, seeking, extinction or drug-induced reinstatement (**Supplementary Figure 3h-l**).

In another set of experiments in male mice, we assessed the effect of Nr4a1 CRISPR knockdown in VP→MDT neurons on reward (**Figure 5a, f; Supplementary Figure 4a-c**). Again, water self-administration was not significantly different between Nr4a1 CRISPR knockdown or LacZ control mice: 2-way RM ANOVA: Session x Virus: F_(7, 182)_= 0.624; p= 0.735 (**Supplementary Figure 4d**). For cocaine self-administration, a 3-way ANOVA showed a significant Day x Drug interaction: Day x Drug: F_(9, 64)_= 3.392; p= 0.001 suggesting that mice developed cocaine self-administration over the 10 days. However a Drug x Virus interaction: Drug x Virus: F_(1, 16)_= 8.054; p= 0.011 further suggested that viral manipulation had an effect of the acquisition of FR1. Tukey’s multiple comparison revealed that LacZ control mice developed cocaine self-administration and were taking more doses compared to their respective saline group for days 6, 9 and 10 (p≤ 0.05 at least). Interestingly, such difference in mice with Nr4a1 CRISPR knockdown was never observed across the 10 days (p>0.999). Moreover, mice with Nr4a1 CRISPR knockdown took significantly less cocaine than their LacZ counterparts for days 8 and 10 (p= 0.009 and 0.002 respectively) (**Figure 5b**). Again, similar results were obtained when assessing active and inactive responses, showing that only the LacZ control group acquired and discriminated the active and inactive ports (**Supplementary Figure 4e**).

**Figure 5.**
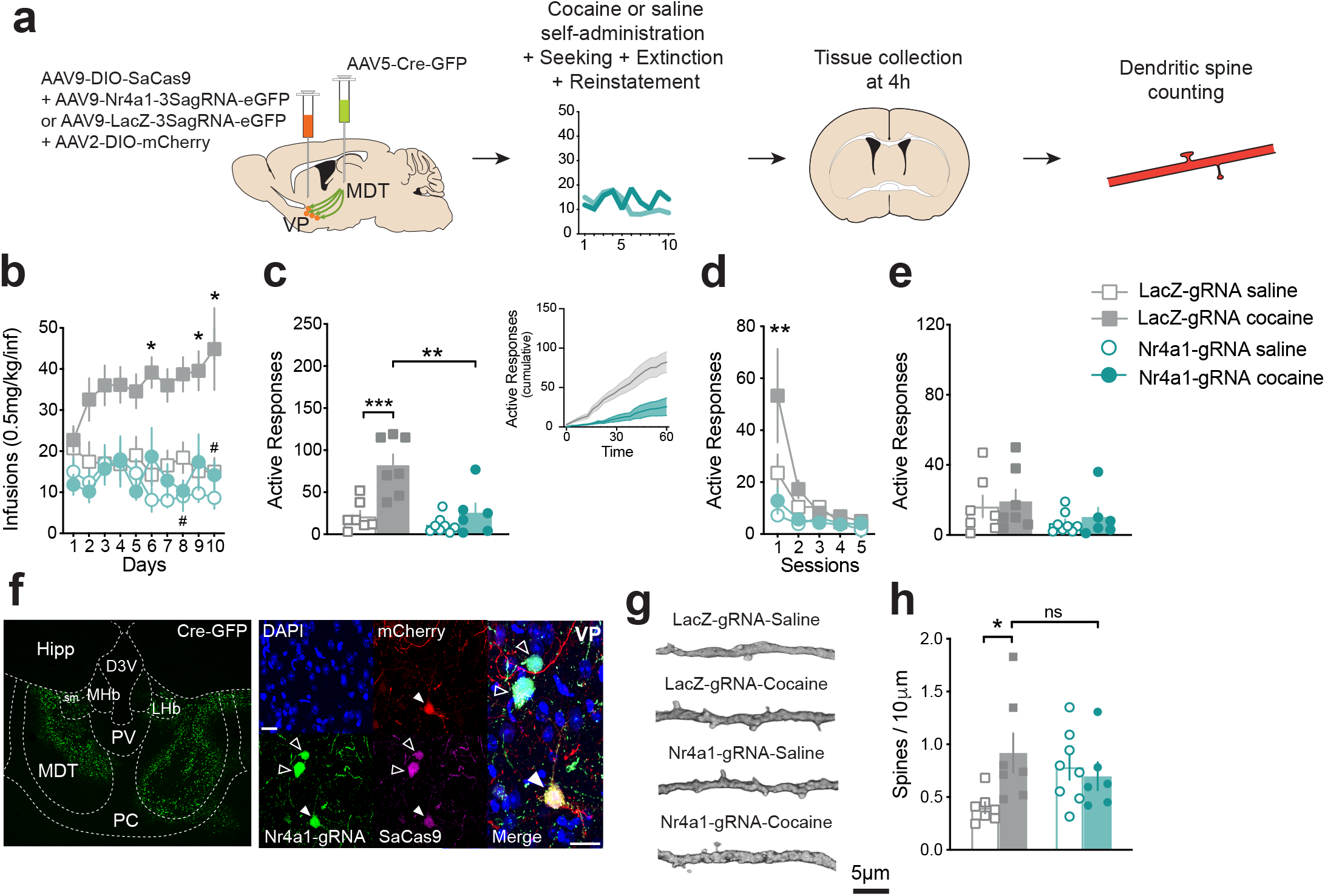
Knockdown of Nr4a1 in VP to MDT neurons blocks FR1 acquisition and alters dendritic spine density. **a**: Schematic of the experimental design: mice received an infusion of retrograde (AAV5) Cre-GFP vector in the MDT and an infusion of DIO-SaCas9 + Nr4a1-sagRNAx3-eGFP (or LacZ-sagRNAx3-eGFP) + DIO-mCherry in the VP prior to 10 days of cocaine (0.5 mg/kg/inf.) or saline self-administration. On day 11, mice were subjected to seeking test (under extinction condition) as well as 5 additional extinction sessions. On day 12, mice were tested for drug- induced reinstatement. Four hours following this last test, tissue was collected for spine density evaluation. n= 7 LacZ-saline, 7 LacZ-cocaine, 8 Nr4a1-gRNA-saline, 6 Nr4a1-gRNA-cocaine; **b**: Mice with Nr4a1 knockdown in VP to MDT neurons did not acquire cocaine self-administration and showed intake levels comparable to saline controls (*p< 0.05 from respective saline, #p< 0.05 from LacZ-cocaine mice); **c**: LacZ-cocaine showed significant seeking behavior compared to their respective saline controls as well as compared to Nr4a1-gRNA-cocaine mice (p< 0.01 and p< 0.001 respectively). Insert represents cumulative active response in cocaine mice during the 1 hr seeking test (5 min bins); **d**: Active responses during extinction test (p< 0.01 from all); **e**: Animals previously taking cocaine were exposed to non-contingent cocaine for drug-induced reinstatement (7.5 mg/kg, i.p.; saline controls received saline, i.p.). None of the groups showed enhanced responding following exposure to a low dose of cocaine; **f**: Left panel: Representative image of MDT injection site for retrograde (AAV5) Cre-GFP viral vector. Right panel: Representative image of cells with Nr4a1 knockdown (DIO-SaCas9 + Nr4a1-gRNA) and sparse DIO-mCherry labelling in the VP. Closed arrowhead: SaCas9+Nr4a1-gRNA+mCherry positive cell, open arrowhead: SaCas9+Nr4a1-gRNA positive cell. Scale bars= 25 µ m; **g**: Representative image of dendritic spines from sparsely labelled VP to MDT neurons from each group. Scale bar= 5 µ m; **h**: Spine density was significantly increased in control mice taking cocaine. Both groups of mice with Nr4a1 knockdown showed elevated spine density levels that did not differ from control mice taking cocaine (p< 0.05, ns: not significant). Hipp: hippocampus, D3V: dorsal 3^rd^ ventricle, MHb: medial habenula, LHb: lateral habenula, sm: stria medularis, PV: paraventricular thalamic nucleus, MDT: mediodorsal thalamus, PC: paracentral thalamic nucleus. See also **Supplementary Figure 4**.

Next, the mice were tested for cocaine seeking behavior. A 2-way ANOVA revealed that both previous drug exposure and viral manipulation had an effect on seeking behavior: Drug x Virus: F_(1, 24)_= 5.770; p= 0.024. Tukey posthoc showed that mice with LacZ gRNA in VP→MDT neurons previously taking cocaine had significantly higher active responses compared to their respective saline controls (p< 0.0007), but also higher responses compared to mice with Nr4a1 CRISPR knockdown previously taking cocaine (p< 0.002) (**Figure 5c**). Following seeking test, mice were evaluated for extinction in similar test conditions. A 3-way ANOVA showed that extinction rate was different depending on the viral manipulation: Session x Virus: F_(4, 12)_= 6.626; p= 0.005.

Tukey posthoc showed that mice with LacZ gRNA in VP→MDT neurons previously taking cocaine had significantly higher active responses compared to their respective saline controls (p= 0.002) and compared to mice with Nr4a1 CRISPR knockdown previously taking cocaine (p= 0.004) for session 1 (**Figure 5d**). Finally, the following day mice were tested for drug-induced reinstatement. Response levels were low in every group and the 2-way ANOVA reported no effect of a low dose of cocaine on active responses: 2-way ANOVA: Drug: F_(1, 24)_= 0.375; p= 0.545 (**Figure 5e**). Mice were perfused four hours following the drug-induced reinstatement test and dendritic spine density was assessed in VP→MDT neurons specifically (**Figure 5g**). A 2- way ANOVA showed a significant Drug x Virus interaction: Drug x Virus: F_(1, 24)_= 5.257; p= 0.030. Tukey posthoc test revealed that spine density was higher in mice with LacZ control previously taking cocaine compared to their respective saline controls (+130.1%; p< 0.05). This was not the case for mice with *Nr4a1* CRISPR knockdown previously taking cocaine (p= 0.966). However, spine density in these mice was increased, although non-significantly (+74.7%; p= 0.43), compared to saline *LacZ* controls. Interestingly, mice with *Nr4a1* CRISPR knockdown previously taking saline did not significantly differ from mice with *LacZ* control previously taking cocaine (p= 0.878) (**Figure 5h**). Moreover, although not significant (p=0.17) spine density in saline mice with *Nr4a1* knockdown was increased compared to LacZ saline controls (+96.7%). Finally, mice with *Nr4a1* CRISPR knockdown previously taking cocaine did not significantly differ from mice with *LacZ* control previously taking cocaine (p= 0.67). Overall, high spine density was seen in mice with *Nr4a1* knockdown regardless of the drug condition further supporting the role of Nr4a1 in dendritic spine dynamics. Spine subtype analysis confirmed this by revealing an overall similar profile for thin spines exclusively (saline: +98.7%; cocaine: +80.4%; both from LacZ saline), suggesting again that changes in *Nr4a1* levels contribute to spinogenesis regulation (**Supplementary Figure 4f**).

## Discussion

In this study we showed that cocaine intake resulted in important transcriptomic alterations in the VP with a major impact on structural plasticity-associated genes. We identified *Nr4a1* as a major gene target and found that its expression was preferentially increased in VP→MDT neurons.

Circuit-specific *Nr4a1* overexpression resulted in enhanced seeking and drug-induced reinstatement while *Nr4a1* knockdown drastically impaired self-administration acquisition. Dendritic spine analyses further revealed that bidirectional *Nr4a1* manipulation markedly altered spinogenesis.

Cocaine exposure alters VP neuronal activity including in specific cell types ^8, 12, 48^. However, the transcriptomic landscape associated with these alterations was largely unexplored. Our gene ontology analysis revealed that a large proportion (55%) of the SDE genes at the “Cellular Component” level were associated with structural plasticity. This suggests that adaptations in connectivity are prominent in VP neurons, consistent with its central role in the reward circuitry ^1^. The subsequent transcriptional regulator analysis of our dataset guided us toward *Nr4a1* (*Nurr77*), an activity-induced transcription factor involved in dendritic spine remodeling ^41^ as well as drug-mediated behaviors ^39, 40^. Interestingly, expression levels of several *Nr4a1* target genes involved in spine remodeling through the Ras/Raf GTPase signaling pathway such as *Plk2, Dlgap2* and *Rgs14* were altered by cocaine. Activation of RAP GTPases supports dendritic spine dynamics ^41^ and bidirectionally regulates cocaine reward in the NAc ^49^. Remarkably, levels of *Nr4a1* and its target genes were not significantly changed in the VP of female mice. Recent work showed that other small GTPases from the Ras superfamily were unaltered by stress and repeated drug intake in the NAc of female mice ^50^. This further suggests that drug exposure may differentially involve small GTPases in males and females, possibly through distinct transcription factor activation patterns ^16^. Our findings suggest that Nr4a1 can mediate both drug intake and cocaine-induced structural plasticity in male mice.

Since the VP is a heterogeneous region with distinct circuits playing differential roles in reward, we assessed *Nr4a1* expression in different VP projection neuron subpopulations. Recent work showed that VP→VTA and VP→LHb neurons either promote or reduce reward-related behaviors ^3, 11^. While these two subpopulations are affected by cocaine ^12^, they did not appear to engage *Nr4a1* signaling. In fact, we found that cocaine intake reduced the proportion of VP→VTA cells expressing *Nr4a1* and that overexpressing *Nr4a1* in this circuit had no effect on cocaine intake or relapse-like behaviors. Since VP→VTA neurons are important for cocaine seeking behaviors ^4^, this further suggests alterations in this circuit likely involve different molecular pathways such as Fos-related signaling ^51, 52^. This could explain why VTA and MDT projecting VP neurons undergo different cocaine-induced synaptic modifications ^12^. Interestingly, the VP→MDT subpopulation was the only neuron population to show both increased proportion of cells expressing *Nr4a1*, and increased Nr4a1 expression levels. Surprisingly, information on the role of this circuit in reward is scarce ^52^, and its involvement in drug-related behaviors is unknown. However, individuals with substance use disorders, drug-induced dopamine release (measured with positron emission tomography) in the MDT correlates with cocaine craving ^53^, and MDT lesions in rats attenuates cocaine intake ^54^. Thus, while both the VP and the MDT are involved in drug addiction, the contribution of the VP→MDT circuit requires deeper characterization. To probe the molecular signature of VP→MDT neurons and their subsequent alteration by cocaine, we conducted a projection-specific evaluation of various cellular markers as well as structural plasticity molecules. Similar to the VP→LHb and VP→VTA neurons, we found that VP→MDT neurons preferentially express parvalbumin ^7^. Consistent with other VP parvalbumin subpopulations ^13^, VP→MDT cells express both *Gad1* and *Vglut2*, supporting both GABAergic and glutamatergic neuron composition ^7, 12^. Additionally, VP→MDT neurons showed high levels of NMDA receptor subunits 1 and 2b as well as GABA receptor subunits α 1 and β 2 suggesting that they receive both glutamatergic and GABAergic inputs. Here, we cannot conclude whether the same VP→MDT neurons express both GABAergic and glutamatergic markers or if they represent two parallel pathways. However, based on findings from other VP circuits, the latter explanation seems the most likely ^13^. We also found that *Chrna4* was highly enriched in these cells, suggesting that these cells could be cholinoceptive ^55^. When assessing VP→MDT neuron gene expression following cocaine self-administration, we extended our *in-situ* hybridization findings of increased *Nr4a1* levels at a 4-h timepoint after drug intake. Moreover, both *Nr4a1* upstream regulator *Mef2c* and downstream target *Plk2* were increased. Plk2’s downstream targets, *Sipa1l1* (SPAR) and *Rap2a*, displayed mRNA enrichment paralleled by reduced *Rasgrf1* mRNA, further supporting the role of this small GTPase signaling in our study and more largely in Nr4a1-mediated adaptations ^41, 44^. This is in line with our findings of increased RAP2 activity after cocaine exposure. Additionally, *Syn1* levels were increased, further corroborating changes in synaptogenesis and neurotransmitter release ^56^. Of note, although basal enrichment of *Penk* and *Tac1* in this cell-population was low, cocaine intake significantly increased levels of both these molecules. Our observations, supported by recent findings ^2, 57^, suggest peptidergic transmission originating from the VP could be involved in motivated behaviors.

To further confirm the role of Nr4a1 in cocaine-related behaviors as well as dendritic spine dynamics we conducted circuit-specific gene manipulation experiments ^58, 59^. With *Nr4a1* overexpression, unrewarded responses during the seeking test were increased (+79%) as well as during drug-induced reinstatement (+47%). This persistent responding is consistent with the role of the MDT in behavioral flexibility and perseverance ^60^. It is likely that our manipulation impaired the transfer of reward value information from the VP to the MDT, leading to inadequate cue information processing and impaired behavioral adjustment ^5, 52, 61^. Indeed, Nr4a1 is involved in spine remodeling mechanisms necessary for synaptic potentiation and memory formation ^41, 62^. Alteration of structural plasticity with *Nr4a1* manipulation may have impaired the adaptation to the changes in action-outcome association, for which VP and MDT are involved ^52, 63^. However, *Nr4a1* manipulations did not impact water self-administration suggesting that the VP→MDT circuit may not be involved in learning action-outcome association for natural rewards but could be engaged in more complex, delayed rewards such as cocaine ^64^. Alternatively, *Nr4a1* manipulation may have altered sensitivity to cocaine. Indeed, *Nr4a1* knockdown in VP→MDT neurons drastically impaired self-administration acquisition. First, this suggests that in addition to its role in early and late cocaine abstinence ^39, 40^, *Nr4a1* also has a role in the development of cocaine intake. Additionally, these results are in line with previous findings in nonhuman primates showing that the MDT is engaged in the early phase of cocaine self-administration acquisition ^65^. Moreover, seminal studies in rats showed that MDT lesion decreases acquisition of cocaine intake, associated with a downward shift in the descending limb of the dose-response curve, suggesting greater sensitivity to cocaine reward ^54^. Given that VP neurons are comparatively more sensitive to intravenous cocaine than NAc or VTA neurons ^8^, changes in VP→MDT structural plasticity could have altered reward sensitivity in our mice. However, the opposite effect on acquisition following *Nr4a1* overexpression was not observed and might have been overshadowed by already high intake levels. Alternatively, the supraphysiological levels of *Nr4a1* achieved with overexpression may have prevented the emergence of circuit adaptations impacting self-administration behavior as it would normally occur with cocaine intake. *Nr4a1* induction with CRISPR activation approaches ^66^ may more closely reproduce physiological gene regulation to potentially highlight differences in FR1 acquisition kinetics. Similarly, we induced *Nr4a1* knockdown prior to cocaine exposure to mimic our overexpression study. This prevented FR1 acquisition and reduced our ability to precisely assess the effect of our manipulation on drug seeking and reinstatement. By blocking the endogenous cocaine-induced activation of *Nr4a1* expression using CRISPR interference ^58, 67^, future studies will allow to more closely reproduce physiological gene regulation and its impact on cocaine-related behaviors.

Alternatively, *Nr4a1* knockdown shortly after self-administration acquisition will provide information on *Nr4a1*-mediated functions after early abstinence. Similarly, gene manipulation after FR1 acquisition will inform on the role of *Nr4a1* on relapse-like behavior following prolonged abstinence ^16^.

Glutamatergic inputs from the subthalamic nucleus, the medial prefrontal cortex (mPFC) and the basolateral amygdala (BLA) increase firing rates in the VP ^7^. This effect is likely enhanced in VP→MDT neurons due to cocaine-induced alterations in glutamatergic receptors on these cells ^12^. *Nr4a1* expression is closely related to excitatory transmission and plays an important role in restoring homeostasis after neuronal overexcitation by regulating dendritic spine density via *Plk2* induction ^39-41, 44, 68^. *Plk2* is a master regulator of GTPases that promote spine pruning, and is thus an important player in synaptic scaling, which involves an intricate interplay between pro- and anti-spinogenesis molecules ^41, 44, 69^. Consistent with this idea, we found that cocaine intake increased *Plk2* expression in bulk VP, and in VP→MDT neurons. Thus, our *Nr4a1* overexpression approach may have partially mimicked cocaine-induced neuronal overactivation, causing blunted cocaine-induced spinogensis and enhanced relapse-like behaviors. Conversely, CRISPR-mediated *Nr4a1* knockdown likely heightened the spinogenesis set point prior to self-administration regardless of the drug condition. We postulate that *Nr4a1* knockdown partially prevented *Plk2* induction, thereby reducing spine pruning, altering the balance of pro- and anti- spinogensis signaling, resulting in a net increase in dendritic spines on VP→MDT neurons. This idea is supported by an almost doubled thin spine density in saline *Nr4a1* knockdown mice. Further, previous studies demonstrated that *Nr4a1* knockdown increases spine density even after basal neuronal activity and its knockdown impairs long-term-potentiation by reducing synaptic strength ^41, 62, 70^. Thus, in our study, *Nr4a1* knockdown may have impaired synaptic potentiation onto VP→MDT neurons, thereby impairing FR1 acquisition. Future studies using input- and output-specific electrophysiological recordings following *Nr4a1* manipulation will increase our understanding of how different VP networks drive acquisition and relapse to cocaine seeking. This will be especially important since several brain regions in key circuits (VP→MDT→mPFC/orbitofrontal cortex (OFC)→NAc→VP or VP→BLA→NAc→VP) exhibit divergent structural changes after cocaine, including spine density increases in mPFC, NAc, and BLA, but spine density decreases in OFC ^1, 71-74^. Moreover, while Nr4a1 closely relates to glutamatergic function, an important role for dopamine receptors in Nr4a1 induction has been reported, especially in striatal GABAergic neurons (see ^39^ for review). Future work evaluating how inhibitory or modulatory synapses are affected by Nr4a1-related spine dynamics will provide a more complete understanding of its role on circuit activity following cocaine intake.

Overall, VP→MDT neurons displayed homeostatic spine remodeling mechanisms induced by *Nr4a1* resulting in circuit adaptations facilitating pro-reward behaviors. While this seems contradictory to the protective role of *Nr4a1* against cocaine-related behaviors described in the NAc ^39, 40^ our work suggests that depending on the brain region, cell-type or circuit, homeostatic mechanisms at the cellular levels can lead to either advantageous or deleterious behavioral outcomes.

In summary, our study uncovers a role for *Nr4a1* in the VP, and specifically VP→MDT neurons, in mediating addiction-like behaviors. We found that Nr4a1 negatively regulates spine dynamics and promotes pro-reward behaviors. Thus, in addition to provide further understanding on the role of *Nr4a1* in cocaine-related behaviors, our work highlights the crucial role of the VP→MDT circuit in substance use disorders.

## Acknowledgments

This work was funded by NIH grants R01MH106500, R01DA038613, R01DA047843 and Israel-US Binational Science Foundation 201725 (to MKL), K99DA050575 (to MEF), F31DA052967 (to EC), T32DK098107 and F32DA052966 (to CAC). Data sharing and visualization via gEAR was supported by grants R24MH114815 and R01DC019370. SST and RJH were supported by the University of Maryland Scholars Program, an initiative of the University of Maryland: MPowering the State. The authors would like to thank Yang Song, Ronna Hertzano and the members of the UMSOM Institute for Genome Science (IGS) as well as Katherine Duarte for their technical help. We also thank Dr. Kristen Maynard and Dr. Keri Martinowich for their help with RNAscope protocols.

## Author contributions

ME and MKL designed the experiments. ME, MEF, HQ, SST and RJH conducted behavioral experiments, MDT and VMR provided animal support. ME and HQ performed RAP2 assay. ME, EYC, RJH and VMR extracted RNA and/or performed qRT-PCR experiments. HN and ME performed *in situ* hybridization. MEF conducted cell-type specific RNA extraction. SST and ME performed *in situ* hybridization quantification. RC, EYC and CAC designed viral constructs. ME performed bioinformatic analyses. ME and MEF conducted neuronal morphology experiments. ME and MKL wrote the manuscript with contributions from all authors.

The authors declare that they have no conflict of interest.

