## Supplementary Figure 1 for "Transcriptome profiling of the ventral pallidum reveals a role for pallido-thalamic neurons in cocaine reward"

**a**

| Category | GO term | # genes | Freq. in set | Corr. p value |
|---|---|---|---|---|
| Biological Process | Signal transduction (GO:0007165) | 63 | 20.3% | p< 4.7844E-6 |
| | Cell communication (GO:0007154) | 50 | 16.1% | p< 2.3360E-11 |
| | Synaptic transmission (GO:0007268) | 34 | 10.9% | p< 5.1779E-12 |
| | Cell projection organization (GO:0030030) | 30 | 9.6% | p< 5.0450E-4 |
| | Cytoskeleton organization (GO:0007010) | 29 | 9.3% | p< 1.0077E-3 |
| | Cell morphogenesis (GO:0000902) | 26 | 8.4% | p< 3.2127E-3 |
| | Reg. of neuron proj. development (GO:0010975) | 25 | 8% | p< 4.7844E-6 |
| | Neuron projection morphogenesis (GO:0048812) | 18 | 5.8% | p< 8.1937E-4 |
| | Actin cytoskeleton organization (GO:0030036) | 17 | 5.5% | p< 2.5829E-3 |
| | Regulation of synaptic plasticity (GO:0048167) | 15 | 4.8% | p< 5.2424E-4 |
| Cellular Component | Cell projection (GO:0042995) | 82 | 25.7% | p< 1.9094E-14 |
| | Neuron projection (GO:0043005) | 68 | 21.3% | p< 1.4402E-19 |
| | Synapse (GO:0045202) | 62 | 19.4% | p< 1.2420E-20 |
| | Cytoskeleton (GO:0005856) | 56 | 17.6% | p< 3.4743E-5 |
| | Synapse part (GO:0044456) | 44 | 13.8% | p< 6.9172E-16 |
| | Dendrite (GO:0030425) | 43 | 13.5% | p< 1.5502E-15 |
| | Postsynaptic membrane (GO:0045211) | 21 | 6.6% | p< 1.3784E-9 |
| | Postsynaptic density (GO:0014069) | 21 | 6.6% | p< 8.2744E-8 |
| | Neuron spine (GO:0044309) | 16 | 5% | p< 1.0984E-7 |
| | Dendritic spine (GO:0043197) | 15 | 4.7% | p< 4.6394E-7 |
| Molecular Function | Cytoskeletal protein binding (GO:0008092) | 31 | 10% | p< 1.8242E-4 |
| | Transmembr. transporter activity (GO:0022857) | 31 | 10% | p< 2.8033E-4 |
| | Kinase activity (GO:0016301) | 22 | 7.1% | p< 8.3849E-3 |
| | Transcription factor activity (GO:0003700) | 21 | 6.8% | p< 1.5338E-2 |
| | Actin binding (GO:0003779) | 15 | 4.8% | p< 8.9181E-3 |

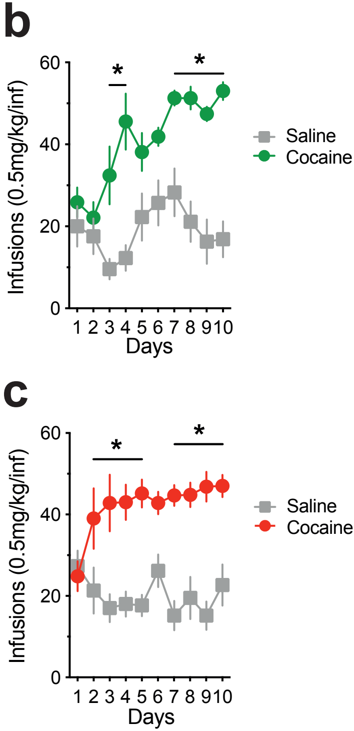


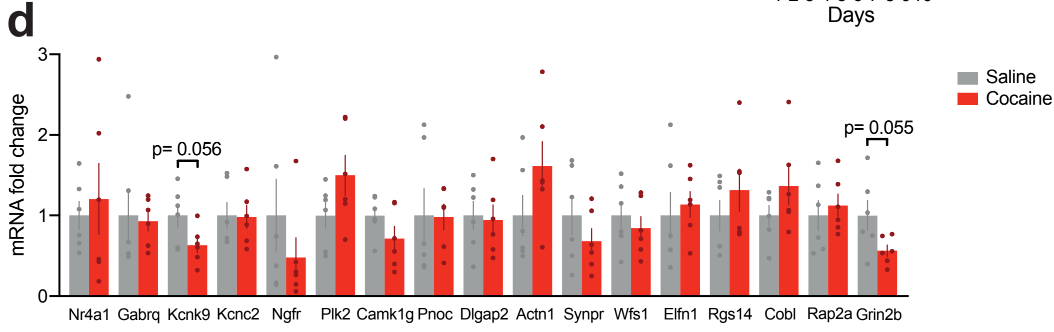
