## Supplementary figures and images for "Transcriptome profiling of the ventral pallidum reveals a role for pallido-thalamic neurons in cocaine reward"

### Supplementary Figure 2

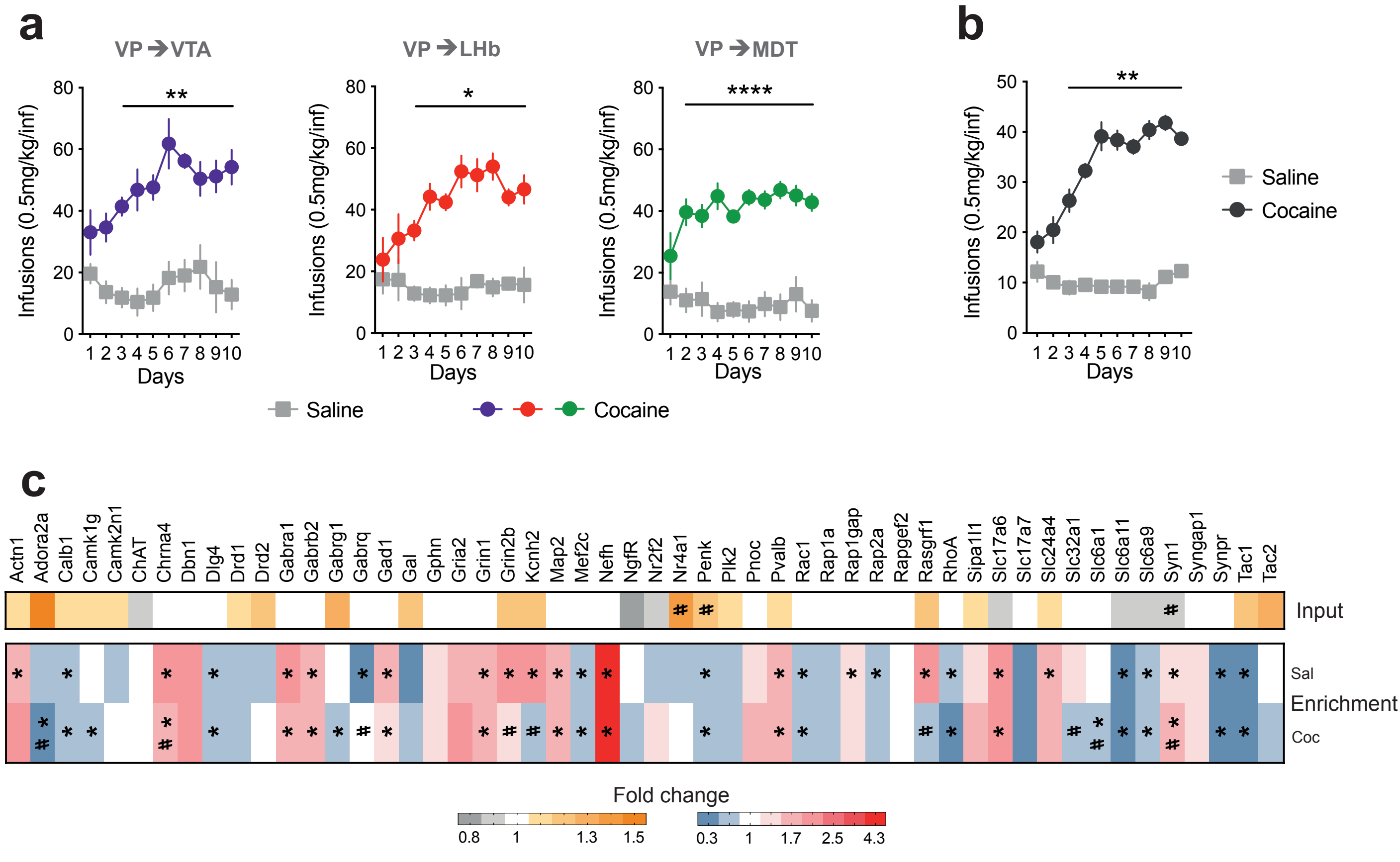

### Supplementary Figure 3

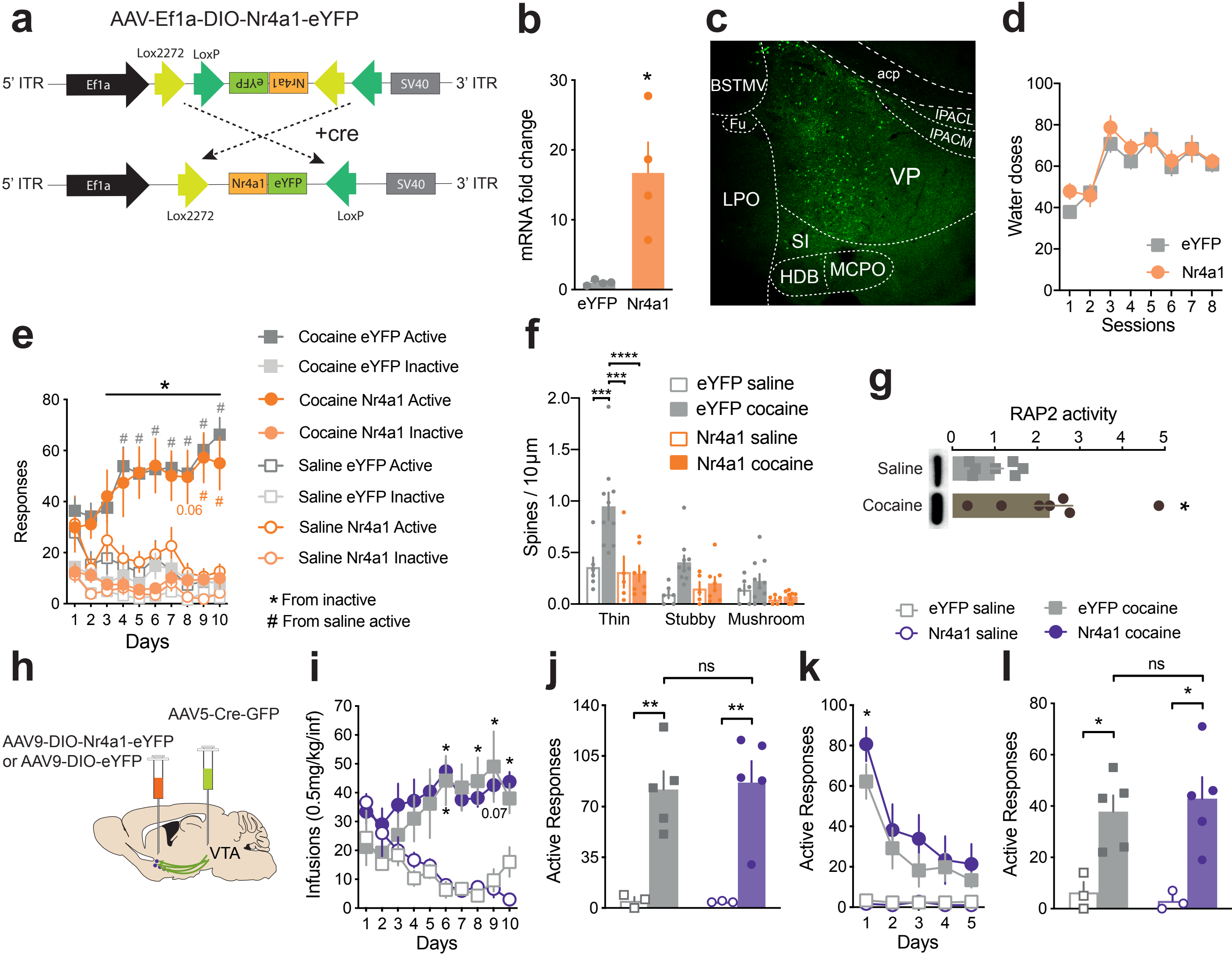

### Supplementary Figure 4

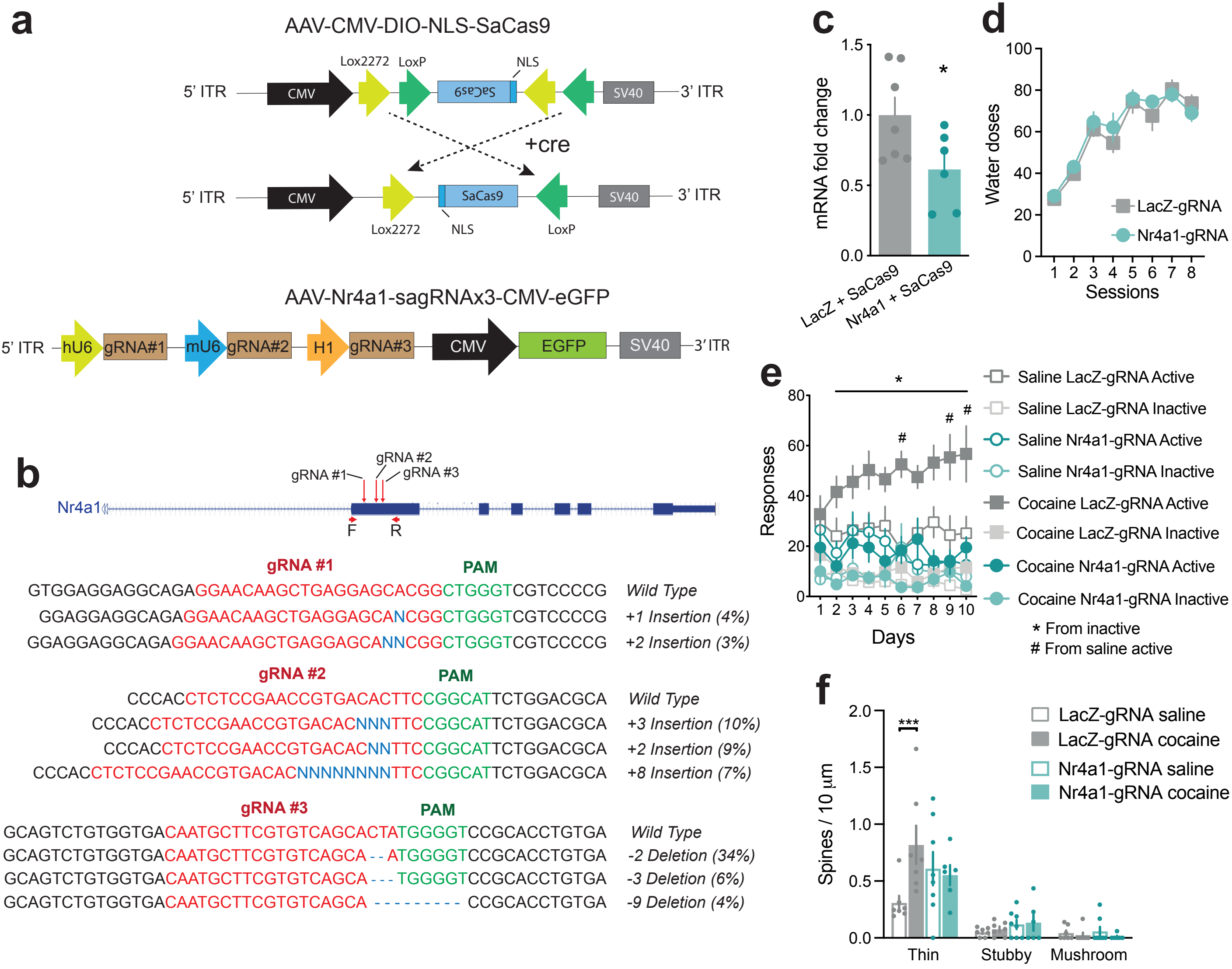
