## Supplementary Methods for "Transcriptome profiling of the ventral pallidum reveals a role for pallido-thalamic neurons in cocaine reward"

### Address Correspondence to

Or

Michel Engeln
Institut de Neuroscience Cognitives et Intégratives d'Aquitaine - UMR 5287
146 rue Léo Saignat
Zone Nord - Bat 1A - 3e étage
33076 Bordeaux cedex
France

ORCID: 0000-0003-3970-7022

Funding: This work was funded by NIH grants R01MH106500, R01DA038613, R01DA047843 and Israel-US Binational Science Foundation 201725 (to MKL), K99DA050575 (to MEF), F31DA052967 (to EC), T32DK098107 and F32DA052966 (to CAC).

Conflict of Interest: The authors declare they have no conflicts of interest.

Supplementary Methods

*Rap2 activity assay*

Rap2 GTPase activity was assessed from whole VP tissue of mice 24 hours after receiving 10 daily cocaine (20mg/kg in saline) or saline intraperitoneal injections. Non-contingent injections were used here because of the number of animals per condition (2x14 mice). Although we did not use contingent drug intake for the assay, the cocaine dose was chosen based on our previous self-administration work in mice [1, 2]. Tissue from 2 mice was pooled per sample and 7 samples were analyzed per condition. Following tissue homogenization, Rap2 activity was measured using a Rap2 activation assay kit following manufacturer's instructions (Cell Biolabs Inc, STA-406-2).

*Nr4a1 CRISPR/Cas9 viral constructs*

The construct for CRISPR-based knockdown of Nr4a1 was designed by Vector Builder. In brief, exons and splice isoforms of Nr4a1 sequence gene was found on the UCSC genome browser database (http://genome.ucsc.edu/). Based on best transcript alignment, gRNA and PAM sequences were design from the 5' exonic codon with benchling.com. We chose multiplexed sgRNA driven by distinct U6 promoters for an efficient knockdown of the target gene. The following sequences were used for the gRNAs: #1gRNA- ccgtgctcctcagcttgttcc; #2gRNA- ctctccgaaccgtgacacttc; #3gRNA- caatgcttcgtgtcagcacta. The multiplexed LacZ gene target was used as control construct. eGFP fluorescence protein expression cassette was added in same vector backbone in under CMV promoter (**Supplementary Figure 4a**).

Indel assay was used to evaluate the capability of our CRISPR construct to induce INsertions or DELetions of nucleotide into the sequence of DNA coding for Nr4a1. Briefly, VP tissue was injected with a mixture of AAV-CMV-DIO-SaCas9, AAV-Nr4a1-sagRNAx3-CMV-eGFP (or AAV-LacZ-sagRNAx3-CMV-eGFP for control) and AAV.hSyn.HI.eGFP-Cre.WPRE.SV40. After 3-4 weeks, genomic DNA was isolated from each sample using 'QuickExtract DNA Extraction Solution' (LucigenQE0905T). In brief, VP tissue was mixed and vortexed (15 sec) with 400μl of QuickExtract Solution. Tubes were transferred to a heat block for 10 minutes at 65°C and vortexed again for 15 seconds. To deactivate the enzymes, the tube was re-heated at 98°C for 2 minutes. 2μl of the sample were used for PCR using Q5 High-Fidelity DNA Polymerase (NEB #M0494S). PCR amplification surrounding each target site from genomic DNA was performed using the following set of primers: Forw: TGTATTCAAGCTCAATATGG and Rev: AGCTTTCTGCAGCCCTAGAGAG. After confirmation of a single PCR product band on the gel, we purified the PCR product using a PCR purification kit (Qiagen #28106). Set of forward and reverse internal primers were used for sanger sequencing at Genomics Core Facility (Formerly Biopolymer Core Facility) at UMB: Sanger Sequence primers: Forw: AGCAACGAGCCCAGGACC, Rev: GTCCTGCAGGACAGAACCAG. CRISPR indel analysis was performed with ICE (Inference of CRISPR Edits; Synthego) tool from synthego.com using .abi files (LacZ .abi file was used as a control). Our sample was verified for CRISPR indels by various ICE tool indel frequency, or the percentage of the cell population that had insertions or deletions (**Supplementary Figure 4b**).

*Statistical analyses*

Samples were excluded if animals did not acquire cocaine self-administration, if not detected (molecular analysis), had inappropriate viral placement (i.e., no/unilateral expression, off-target Cre injection), or failed Grubbs' outlier test. This corresponds to the exclusion of: 2 saline mice in the VP→VTA group, 3 saline and 2 cocaine mice in the VP→LHb group and 2 saline mice in the VP→MDT group of our *in situ* hybridization experiment due to off-target Cre-injection or loss of catheter patency. In the Nr4a1 overexpression experiment: 2 eYFP-saline, 3 Nr4a1-saline, 1 eYFP-cocaine and 3 Nr4a1-cocaine mice due to off-target Cre-injection, no/unilateral expression or loss of catheter patency. In the Nr4a1 knockdown experiment: 2 saline LacZ-gRNA, 1 saline Nr4a1-gRNA, 1 cocaine LacZ-gRNA and 2 cocaine Nr4a1-gRNA mice due to off-target Cre-injection, no/unilateral expression or loss of catheter patency.
