## Supplementary Material 1 for "Transcriptome profiling of the ventral pallidum reveals a role for pallido-thalamic neurons in cocaine reward"

Supplementary Figure 1

**a**: Table summarizing the significant GO terms at the Biological process, Cellular Component and Molecular function obtained following RNA sequencing in male mice. Number of significantly differentially expressed genes from our dataset belonging to each GO term, their frequency in our dataset, as well as their corrected p value are displayed; **b**: Male mice used to validate RNA sequencing data rapidly developed self-administration and showed significantly higher numbers of cocaine infusions compared to Saline for day 3 and 4 as well as from day 7 onward: 2-way RM ANOVA: Day x Drug: $F_{(9, 108)}= 4.529$; $p< 0.0001$; Sidak posthoc: $p< 0.01$ at least; **c**: Female mice self-administering cocaine showed high levels of intake compared to their saline counterparts from day 2 to day 5 and from day 7 onward: 2-way RM ANOVA: Day x Drug: $F_{(9, 90)}= 3.746$; $p= 0.0005$; Sidak posthoc: $p< 0.05$ at least; **d**: Tissue from the VP of female mice was collected 24h hours following the last self-administration session. RNA levels were quantified for Nr4a1 and its target genes (found in our RNA sequencing analysis done in male mice). None of the genes showed significant changes in females: Nr4a1: $t_{10}= 1.106$, $p= 0.294$; Gabrq: $t_{10}= 0.207$, $p= 0.839$; Kcnk9: $t_{10}= 2.153$, $p= 0.056$; Kcnc2: $t_{10}= 0.076$, $p= 0.940$; NgfR: $t_{10}= 1.016$, $p= 0.333$; Plk2: $t_{10}= 1.684$, $p= 0.123$; Camk1g: $t_{10}= 1.553$, $p= 0.151$; Pnoc: $t_{10}= 0.045$, $p= 0.964$; Dlgap2: $t_{10}= 0.213$, $p= 0.835$; Actn1: $t_{10}= 1.548$, $p= 0.152$; Synpr: $t_{10}= 1.101$, $p= 0.296$; Wfs1: $t_{10}= 0.0704$, $p= 0.497$; Elfn1: $t_{10}= 0.0415$, $p= 0.686$; Rgs14: $t_{10}= 0.970$, $p= 0.354$; Cobl: $t_{10}= 1.370$, $p= 0.200$. Some genes showing significance in VP→MDT neurons of males were also evaluated in bulk VP from female mice but showed no significant changes: Rap2a: $t_{10}= 0.527$, $p= 0.609$; Grin2b: $t_{10}= 2.176$, $p= 0.055$.

Supplementary Figure 2

**a**: Male mice received an AAV5-Cre infusion in one of the 3 target structures (VTA, LHb or MDT) in order to label specific VP projection neurons. They were then subjected to self-administration. Each group quickly developed self-administration and showed significantly higher numbers of cocaine infusions compared to saline: VTA: 2-way RM ANOVA: Day x Drug: $F_{(9, 72)}= 2.506$; $p= 0.014$; Sidak posthoc: $p< 0.01$ at least from day 3 onward; LHb: 2-way RM ANOVA: Day x Drug: $F_{(9, 72)}= 3.977$; $p= 0.0004$; Sidak posthoc: $p< 0.05$ at least from day 3 onward; MDT: 2-way RM ANOVA: Day x Drug: $F_{(9, 72)}= 3.124$; $p= 0.003$; Sidak posthoc: $p< 0.0001$ from day 2 onward. *$p< 0.05$, **$p< 0.01$, ****$p< 0.0001$ from saline group; **b**: RiboTag male mice received an AAV5-Cre infusion in the MDT in order to label ribosomes of VP neurons specifically projecting to this structure. 27 mice were subjected to saline and 26 mice were subjected to cocaine self-administration. Self-administration data from each group were pooled (n= 4-5 mice per sample; n= 6 samples per drug condition) in order to conduct subsequent gene expression analysis. Mice showed significantly higher numbers of cocaine infusions compared to saline: 2-way RM ANOVA: Day x Drug: $F_{(9, 90)}$= 19.48; p< 0.0001; Sidak posthoc: p< 0.01 at least from day 3 onward. **p< 0.01 from saline group; **c**: VP tissue was collected 4h following the last self-administration session. Gene expression analysis from VP → MDT neurons was performed using NanoString on 6 samples per drug condition (n= 4-5 mice pooled per sample). Input (i.e., non-immunoprecipitated RNA) was compared between mice taking saline and mice taking cocaine (top row). Gene enrichment (immunoprecipitated vs. input) in VP → MDT neurons was analyzed for each drug condition (bottom rows). *p< 0.05 from input; #p< 0.05 from saline. Complete statistics are available in **Suppl Table 3**.

Supplementary Figure 3

**a**: Schematic of DIO-Nr4a1-eYFP vector; **b**: Nr4a1 overexpression in the VP using AAV9-DIO-Nr4a1-eYFP (+ AAV5-Cre-GFP) significantly increased Nr4a1 levels (+1,574%) compared to AAV9-DIO-eYFP (+ AAV5-Cre-GFP) controls: $t_6$ = 3.606; p= 0.011. *p< 0.05 compared to eYFP controls; **c**: Representative image of Nr4a1 overexpression in VP→MDT cells: VP: ventral pallidum, acp: anterior commissure, post limb, IPACL: interstitial nu. of post limb of ac. lateral, IPACM: interstitial nu. of post limb of ac. medial, SI: substantia innominata, HBD: nu. of horizontal limb of diagonal band, MCPO: magnocellular preoptic nucleus, BSTMV: bed nucleus of stria terminalis medioventral, Fu: bed nucleus of stria terminalis fusiform, LPO: lateral preoptic area; **d**: Male mice receiving Nr4a1 overexpression in VP→MDT neurons showed no difference in natural reward (water) self-administration compared to eYFP controls: 2-way RM ANOVA: Session x Virus: $F_{(7, 189)}$= 0.799; p= 0.589; **e**: Mice from both cocaine groups made significantly more active responses than inactive responses from day 3 onward and significantly more active responses compared their respective saline groups starting at day 4 (eYFP) and Day 9 (Nr4a1): 3-way ANOVA (eYFP): Day x Drug x Lever: $F_{(9, 126)}$= 4.566; p< 0.0001,Tukey posthoc: p< 0.05; 3-way ANOVA (Nr4a1): Day x Drug x Lever: $F_{(9, 99)}$= 2.123; p= 0.034, Tukey posthoc: p< 0.05; **f**: Cocaine intake increased thin spines density in eYFP control mice exclusively: 3-way ANOVA: Virus x

Drug: $F_{(1, 27)}$= 10.22; p= 0.003, Tukey posthoc: p< 0.001 from eYFP saline and Nr4a1 saline, p< 0.0001 from Nr4a1 cocaine; **g**: Rap2 activity assay was conducted 24 hours following 10 days of cocaine (20/mg/kg, i.p. in saline) or saline exposure. Rap2 activity was increased in the VP of mice exposed to cocaine (p<0.05). n= 7/group; **h**: Schematic of the experimental design: Male mice received an infusion of retrograde (AAV5) Cre-GFP vector in the VTA and an infusion of DIO-Nr4a1-eYFP (or DIO-eYFP) + DIO-mCherry in the VP prior to 10 days of cocaine (0.5 mg/kg/inf.) or saline self-administration; **i**: Both groups show significantly higher intake compared to their respective controls: 3-way ANOVA: Day x Drug: $F_{(9, 63)}$= 7.271; p< 0.0001, Tukey posthoc: p< 0.05 at least for day 6, 8, 9 for eYFP and day 6, 10 for Nr4a1 from respective saline. There was no effect of the virus: 3-way ANOVA: Virus x Drug: $F_{(1, 7)}$= 0.070; p= 0.798; **j**: Both groups showed significant seeking behavior compared to their respective saline controls: 2-way ANOVA: Drug: : $F_{(1, 12)}$= 35.79; p< 0.0001, Tukey posthoc: p< 0.01. Both cocaine groups were not significantly different from each other: 2-way ANOVA: Drug x Virus : $F_{(1, 12)}$= 0.051; p= 0.824, Tukey posthoc: p= 0.986; **k**: Active responses during extinction test. Both groups show similar rate of extinction: 3-way ANOVA: Session x Drug: $F_{(4, 32)}$= 14.05; p< 0.0001, Tukey posthoc: p< 0.05 from respective saline controls for session 1; **l**: Animals previously taking cocaine were exposed to non-contingent cocaine for drug-induced reinstatement (7.5 mg/kg, i.p.; saline controls received saline, i.p.). Both eYFP-cocaine and Nr4a1-cocaine mice made significantly more active responses compared to their respective control: 2-way ANOVA: Drug: $F_{(1, 12)}$= 23.88; p= 0.0004, Tukey posthoc: p< 0.05. However, active responses were not significantly different between both cocaine groups: 2-way ANOVA: Virus: $F_{(1, 12)}$= 0.013; p= 0.910, Tukey posthoc: p= 0.942.

Supplementary Figure 4

**a**: Schematic of DIO-SaCas9 and Nr4a1 gRNA viral constructs; **b**: Our Nr4a1 CRISPR vector contained 3 guide RNAs (gRNA) targeted at 3 different loci of the Nr4a1 gene. For each gRNA, the most frequent number of insertions/deletion in the sequenced population is represented in %. PAM: protospacer adjacent motif; **c**: Nr4a1 knockdown in the VP using

AAV9-Nr4a1-SagRNAx3-eGFP (+ DIO-SaCas9 + AAV5-Cre-GFP) significantly decreased Nr4a1 levels (-38.6%) compared to AAV9-LacZ-SagRNAx3-eGFP (+ DIO-SaCas9 + AAV5-Cre-GFP) controls: $t_{11}$= 2.266; p= 0.044. *p< 0.05 compared to LacZ controls; **d**: Male mice with Nr4a1 knockdown in VP→MDT neurons showed no difference in natural reward (water) self-administration compared to LacZ controls: 2-way RM ANOVA: Session x Virus: $F_{(7, 182)}$= 0.624; p= 0.735; **e**: Only mice from the LacZ cocaine groups made significantly more active responses than inactive responses from day 2 onward and significantly more active responses compared their respective saline groups for day 6, 9, 10: 3-way ANOVA: Drug x Lever: $F_{(1, 12)}$= 7.269; p= 0.019,Tukey posthoc: p< 0.05; **f**: Cocaine intake significantly increased thin spines density in LacZ control mice: 3-way ANOVA: Spine type x Virus x Drug: $F_{(2, 33)}$= 5.257; p= 0.010, Tukey posthoc: p= 0.0002 from LacZ saline.

Supplementary Table 1

Complete gene list from our RNA sequencing of VP following self-administration. FC: fold change; LFC: log fold change; FDR: false discovery rate.

Supplementary Table 2

Primer sequences. First tab: list of primer sequences used for qRT-PCR. Second tab: list of primer sequences used for amplification before NanoString assay. Third tab: list of target sequences used for the NanoString assay.

Supplementary Table 3

Detailed statistics for the NanoString assay. Bold font highlights significant p values (<0.05 at least).
